# Regulatory interplay between RNase III and asRNAs in *E. coli;* the case of AsflhD and the master regulator of motility, *flhDC*

**DOI:** 10.1101/2021.05.11.443715

**Authors:** Maxence Lejars, Joël Caillet, Maude Guillier, Jacqueline Plumbridge, Eliane Hajnsdorf

## Abstract

In order to respond to ever-changing environmental cues, bacteria have evolved resilient regulatory mechanisms controlling gene expression. At the post-transcriptional level, this is achieved by a combination of RNA-binding proteins, such as ribonucleases (RNases) and RNA chaperones, and regulatory RNAs including antisense RNAs (asRNAs). AsRNAs bound to their complementary mRNA are primary targets for the double-strand-specific endoribonuclease, RNase III. By comparing primary and processed transcripts in an *rnc* strain, mutated for RNase III, and its isogenic wild type strain, we detected several asRNAs. We confirmed the existence of RNase III-sensitive asRNA for *crp*, *ompR*, *phoP* and *flhD* genes, encoding master regulators of gene expression. AsflhD, the asRNA to the master regulator of motility *flhDC*, is slightly induced under heat-shock conditions in a sigma24 (RpoE)-dependent manner. We demonstrate that expression of AsflhD asRNA is involved in the transcriptional attenuation of *flhD* and thus participates in the control of the whole motility cascade. This study demonstrates that AsflhD and RNase III are additional players in the complex regulation ensuring a tight control of flagella synthesis and motility.

**Importance:** The importance of asRNAs in the regulation of gene expression has long been underestimated. Here, we confirm that asRNAs can be part of layered regulatory networks since some are found opposite to genes encoding global regulators. In particular, we show how an antisense RNA (AsflhD) to the gene expressing a transcription factor serving as the primary regulator of bacterial swimming motility (FlhD_4_C_2_) is involved in the transcriptional attenuation of *flhD*, which in turn impacts the expression of other genes of the motility cascade. The role of AsflhD highlights the importance of discrete fine-tuning mechanisms in the control of complex regulatory networks.

## Introduction

In eukaryotes, hundreds of RNA-binding proteins (RBPs) and multiple classes of regulatory RNAs are involved in the complex regulation of gene expression (splicing, editing…). In bacteria, the relative scarcity of RBPs and regulatory RNAs, led to the supposition that they provided only accessory contributions to the major bacterial gene regulatory mechanisms. An important obstacle in deciphering regulatory networks is their multi-component nature and the existence of “missing links” between regulators and their targets. These intermediates can have both positive and negative impacts on gene expression, leading to compensatory effects upon removal of one of them (1–6). Hence, the fine-tuning of gene expression is far from fully understood in many if not most cases.

Many bacterial small RNAs (sRNAs) are regulators that base-pair with RNA, with their genes located in *trans* to their targets and acting by short, imperfect regions of base-pairing. This property allows them to act on multiple targets. In contrast, the genes for antisense RNAs (asRNAs) are located in *cis* to their complementary target and thus have, in most cases, a single dedicated target. Fewer asRNAs have been described as compared to sRNAs probably because of their high lability, their low conservation among species and because they were usually considered the products of pervasive transcription arising from leaky terminators (7–10).

Initially, asRNAs were identified on mobile genetic elements (prophages and plasmids), with their only purpose to control their replication and partition. The importance of asRNAs was later demonstrated to extend to almost all kinds of biological processes (11), as in the case of type I toxin-antitoxin systems, involved in persistence, in which the toxin mRNA is neutralized by an asRNA that induces degradation and/or translation inhibition (12). Furthermore, the double-strand-specific RNase III has been known to be an important player in asRNA regulation, as in the case of the regulation of plasmid copy number and toxin-antitoxin systems (13–14).

The mechanisms of action of asRNAs are diverse. They can negatively regulate transcription by interference due to the collision of two converging RNA polymerases or by attenuation due, in some cases, to the stabilization of a terminator structure in the mRNA upon binding of the asRNA (15, 16). However, despite complete complementarity, the interaction of asRNA and its target requires, in some cases, formation of an intermediate called “kissing complex” (13, 17). These interactions can have negative or positive consequences on gene expression since they induce modifications to the RNA secondary structure and/or physically interfere with the activity of other regulators (18, 19). Very often, the mechanism by which a specific asRNA regulates its target remains unclear due to the impossibility to modify the sequence of the asRNA independently of its target.

Various approaches have been used to enrich the *E. coli* transcriptome for double-stranded RNAs, which are the presumed intermediates in asRNA regulation. Studies using inhibition of Rho-dependent transcription termination demonstrated that pervasive antisense transcription is common in almost all loci in *E. coli* (20). In 2010, one thousand asRNAs were identified suggesting their importance in the control of gene expression (9). Another study focusing on the mapping of transcriptional units highlighted the presence of 498 asRNAs mostly from overlapping untranslated regions (UTRs) within convergent or divergent operons (21). Immunoprecipitation of double-stranded RNAs using specific antibodies allowed the identification of 200 asRNAs of which 21 were validated as RNase III-degraded asRNAs (22). In a fourth study, primary transcripts were isolated by selective tagging allowing the identification of 212 asRNA transcription start sites (asTSSs) (23). More recently, p19 viral protein capture of double-stranded RNAs identified 436 asRNAs (24). Unexpectedly, there is a little overlap between the identified asRNAs from these different studies, which may be due to technical bias, but may also depend on the genetic context and/or the environmental conditions.

While previously published works focused on the identification of asRNAs, we aimed to characterize physiologically relevant asRNAs. Some time ago we performed a transcriptome analysis of an *rnc* mutant compared to its isogenic wt strain. We used a tailored RNA-seq approach described previously (25). In agreement with other published genomics experiments (22–23, 26) several candidate asRNA were detected, which were stabilized in the strain lacking RNase III activity. We were surprised to see that many were asRNA to genes of important regulators which raised the question of whether or not they could have a physiological impact on the expression and function of their target regulator and hence on the downstream regulon. We first confirmed that RNase III modulates the level of 4 of these antisense transcripts and then concentrated on the asRNA to *flhD*. *flhD* is the first gene of the *flhDC* operon encoding the master regulator of swimming motility. We find that AsflhD is involved in the direct repression of the transcription elongation of *flhD,* which provides an additional regulatory layer to the complex cascade of motility in enterobacteria.

## Results

### Characterization of asRNAs stabilized upon RNase III inactivation

An RNA-seq analysis in a wt and its *rnc*105 derivative strain was performed by tagging transcripts according to their 5′-phosphorylation status, allowing to distinguish between 5’-triphosphate fragments (primary transcripts TSS), monophosphate 5’-fragments (processed transcripts, PSS) and internal fragments resulting from the fragmentation (INT) (25). The depth of sequence coverage was not sufficient for a compilation of all asRNAs. Instead we looked manually for antisense reads covering the translation signals of mRNAs that were enriched upon RNase III inactivation. Potential asRNA promoter regions were deduced by examination of the TSS and PSS fractions in the *rnc* strain. The RNase III processing sites in the wild-type were usually not obvious since they presumably provoked the rapid degradation of the asRNA. We selected 4 asRNAs to the *crp*, *ompR*, *phoP* and *flhD* transcripts encoding important global regulators for verification by northern blot (Fig. 1). We note that they had all been proposed as antisense transcripts in one or more of the previous genomic studies (9, 21–24).

**Figure 1:**
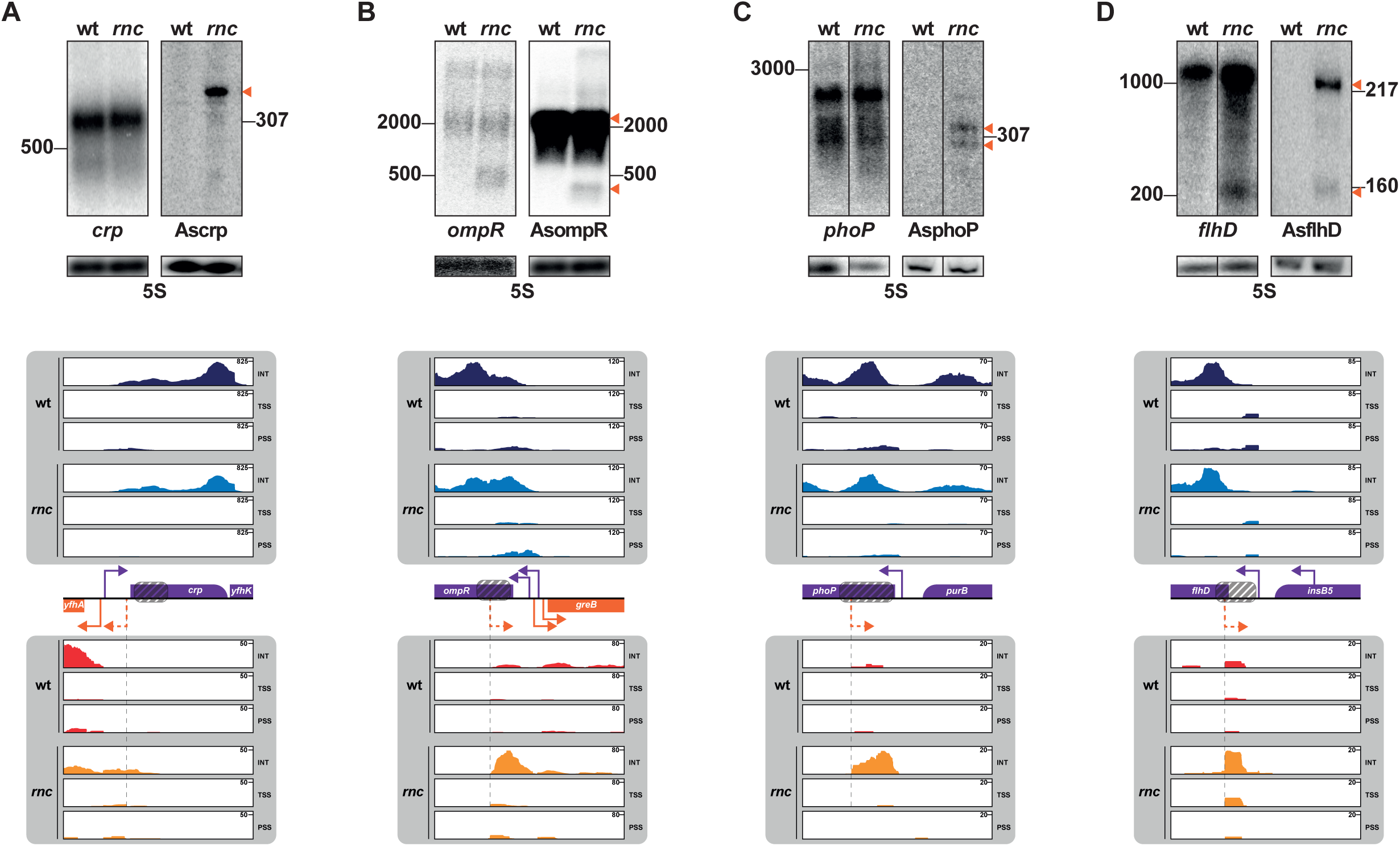
RNase III inactivation stabilizes asRNAs. The RNA-seq reads were aligned to the genome of reference (MG1655, GenBank identifier U00096.3, https://www.ncbi.nlm.nih.gov/nuccore/U00096.3) and visualized with the Integrative Genomic Viewer (IGV) 2.4.2 software (http://software.broadinstitute.org/software/igv/). The fractions isolated during the RNA-seq analysis are color-coded; strand of the target gene in wt (dark blue) and in *rnc* mutant (light blue); strand of the asRNA gene in wt (red) and in *rnc* mutant (orange). Reads corresponding to transcription start site (TSS), processing sites (PSS) and internal fragments (INT) are indicated. The scale for the absolute number of reads identified is indicated on the top right of each lane. The schemes indicate the localization of the ORFs and known promoters (plain bent arrows) and the putative antisense promoters deduced from TSS data (dashed bent arrows). Detection of asRNAs (orange triangle) to *crp* (A), *ompR* (B), *phoP* (C) and *flhD* (D). RNAs extracted from exponentially grown N3433 (wt) and IBPC633 (*rnc*) strains were analyzed on agarose or denaturing acrylamide gels and northern blots were probed by using pairs of complementary uniformly radio-labeled RNA probes to the same region of each target and a primer complementary to the 5S rRNA. A scheme of the probed loci is shown under each panel. The position of the probes relative to the DNA sequence is indicated by a dashed box. The putative asRNA promoter is indicated by a dashed orange bent arrow when it could be predicted from RNA-seq data while known promoters are indicated by plain arrows (purple for genes located on the opposite strand from the detected asRNA). To note, *crp* and AsompR were successively probed on the same membrane, thus they share the same loading control. The membranes shown for *phoP* and AsphoP correspond to zero time points of the stability experiment presented in figure 2-A. It should be noted that the *flhD* mRNA detected corresponds in size to the co-transcript *flhDC* (1200 nts).

The *crp* gene encodes the major regulator of carbon catabolite repression and it was shown previously to be transcriptionally regulated by a transcript initiated from a divergently expressed promoter 3 base-pairs upstream and on the opposite strand (27–28). This transcript, now known to express the *yhfA* gene, was detected in the wt strain (Fig. 1-A). Additional asRNAs, were stabilized in the *rnc* strain in the 3 fractions. The TSS of one species is located 20 nts upstream of the *crp* translation start on the opposite strand (shown by an orange dotted arrow on Fig. 1A) and in addition there is extensive asRNA, complementary to the *crp* coding sequence and 5’-UTR (INT). We performed northern blots using complementary probes hybridizing to positions 13 to 441 of the *crp* ORF to detect both the *crp* mRNA and its antisense transcript. An asRNA, named Ascrp, of about 350 nts accumulates only in the mutant (Fig. 1-A).

The *ompR* gene encodes the response regulator of a two-component system involved in cell wall homeostasis and response to low pH, EnvZ-OmpR (29–32). We observed an asTSS 147 nts downstream from the AUG (Fig. 1-B) in wt and enhanced in the *rnc* mutant. Northern blotting with complementary probes corresponding to the 5’-end of the *ompR* ORF confirmed the presence of the *ompR-envZ* transcript and a long asRNA transcript, AsompR, of about 2000 nts, which is likely to also encode the divergently expressed *greB* gene. In addition, in the mutant, smaller fragments (less than 500 nts) are detected for both *ompR* and AsompR (Fig. 1-B), likely corresponding to a stable duplex between the sense and asRNA transcripts in the absence of RNase III.

The *phoP* gene encodes the response regulator of the PhoQ-PhoP two-component system, involved in cell wall homeostasis and in response to low magnesium (33–34). We observed an asRNA upon RNase III inactivation in the INT fraction about 282 nts downstream from the *phoP* AUG. Northern blot with complementary probes corresponding to the 5’-end of the *phoP* ORF confirmed the accumulation of two fragments (AsphoP) about 300-320 nts long in the mutant (Fig. 1-C).

The *flhDC* genes are co-transcribed and together they encode the master regulator of motility, FlhD_4_C_2_ (35). We detected an asRNA to *flhD* mRNA (AsflhD) accumulating in the mutant, initiated 22 nts downstream from the *flhD* AUG. Northern blot analysis with probes hybridizing to the 5’-UTR of *flhD* confirmed the accumulation of AsflhD, with a major fragment about 220 nts and one minor fragment about 160 nts upon RNase III inactivation. Together, there was an increase in the amount of the full-length *flhD* mRNA in the mutant and the stabilization of a *flhD* fragment of approximate size 220 nts (Fig. 1-D).

All these asRNAs are partially or completely processed by RNase III since they are only visible in the *rnc* strain. Crp, OmpR, PhoP and FlhD are major regulators of gene expression in *E. coli*, all involved in the control of large regulons (RegulonDB v 10.5 (36)). We wondered whether these asRNAs and RNase III have a functional regulatory role and thus affect cell physiology. We studied in more details asRNAs to *phoP* and *flhD*, two regulators tightly controlled at both the transcriptional and post-transcriptional levels.

### Regulation of phoP and AsphoP by RNase III

The RNA-seq profiles suggested that AsphoP may be transcribed from an asTSS located 282 nts downstream from the translation start of *phoP* mRNA. An asRNA derived from this TSS was confirmed by northern blot, probing for the 5’-region of the *phoP* ORF (Fig 1-C). In addition, the *rnc* mutation slightly increases *phoP* stability and amount but induces very large increases in the stability and level of AsphoP (Fig. 2-AB). Candidate consensus -10 and -35 sequences are located just upstream of AsphoP TSS (Fig. 2-C). To validate this potential promoter, we constructed a P_AsphoP_-*lacZ* transcriptional fusion containing 150 nts before and 15 nts after the putative TSS of AsphoP, with the wt sequence (P_AsphoP_^wt^) and also with mutations decreasing the agreement with the consensus in the predicted -35 and -10 boxes (P_AsphoP_^-^) (Fig. 2-D). The mutated AsphoP promoter (P_AsphoP_^-^) strongly decreased the expression of P_AsphoP_-*lacZ* (20-fold), confirming it as the endogenous AsphoP promoter (Fig. 2-E). The activity of the P_AsphoP_^wt^-*lacZ* fusion decreased 2-fold in the mutant implying that RNase III positively regulates AsphoP. In summary, RNase III positively controls the transcription of AsphoP and also participates to the degradation of both *phoP* and AsphoP transcripts.

**Figure 2:**
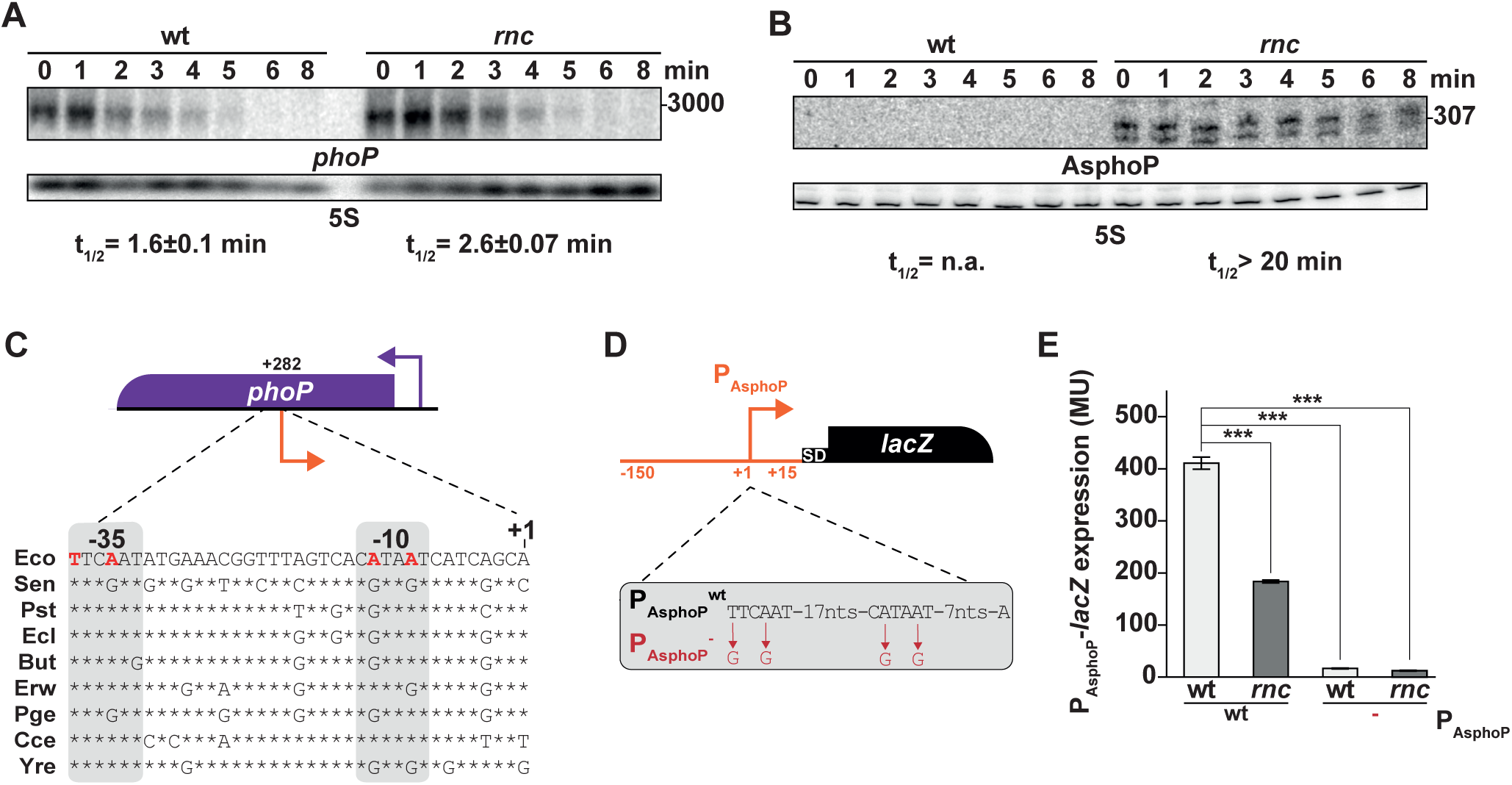
AsphoP and phoP levels are regulated by RNase III. (A) and (B) Stability of *phoP* and AsphoP respectively in the wt (N3433) and the *rnc* (IBPC633) strains. Samples were removed at the times indicated in minutes after rifampicin (500 μg/mL) addition. Half-lives (t_1/2_) were calculated using linear regression and are indicated under each condition. Control of loading was assayed by probing for the 5S rRNA. (C) Genetic structure of the *phoP* locus and alignment of the promoter sequence of AsphoP from selected bacterial species. P_AsphoP_ (orange bent arrow), is positioned relative to the translation start of *phoP* mRNA (+282). Nucleotide sequences correspond to the following genomes, Eco, *Escherichia coli* MG1655 (NC_000913.3), Sen, *Salmonella enterica* LT2 (CP014051.2), Pst, *Pantoea stewartia* ZJ-FGZX1 (CP049115.1), Ecl, *Enterobacter cloacae* NH77 (CP040827.1), But, *Buttiauxella* sp. 3AFRM03 (CP033076.1), Erw, Erwinia sp. J780 (CP046509.1), Pge, Pluralibacter gergoviae (LR699009.1), Cce, *Clostridium cellulovorans* 743B (CP002160.1), Yre, *Yokenella regensburgei* W13 (CP050811.1). Nucleotides in red were mutated to inactivate P_AsphoP_, stars represent conserved nucleotides as compared to Eco. The -35 and -10 motifs of AsphoP are boxed. (D) Genetic structure of the transcriptional AsphoP-*lacZ* reporter fusion. Nucleotides in red indicate the mutations inactivating the AsphoP promoter (P_AsphoP_^-^). (E) Expression of β-galactosidase activity was determined from the P_AsphoP_-*lacZ,* P_AsphoP_*^-^-lacZ* fusions in the wt strain (ML421 and ML422), and in their *rnc* derivatives (ML424 and ML425 respectively). Values are means of three biological replicates for each strain, and error bars are standard deviations. Statistical significance was determined by a heteroscedastic two-tailed t test (*** for p-values ≤0.001).

Sequence comparison with other bacterial species showed that although the region of the AsphoP promoter is moderately well conserved, there are several A to G substitutions in the -10 box at positions -9 and -12, suggesting that this promoter may be inactive in these genomes (Fig. 2-C). This, in turn, implies that, if AsphoP has any function, it could be limited to *E. coli* K-12 and have been counter-selected in these other species or more likely, represents a novel, evolving trait.

### Physiological expression of AsflhD

Figure 1 shows a corresponding increase in the amounts of *flhD* and the appearance of a smaller mRNA fragment in the *rnc* mutant. Intriguingly 40 years ago it was noted that RNase III was involved in the swimming activity of *E. coli* and *rnc* mutants were immotile (37). In this work we have investigated whether RNase III could exert this effect *via* AsflhD.

The RNA-seq profiles revealed an asTSS 22 nts downstream from the translation start of *flhD* as reported by Dornenberg *et al.* (9). A candidate promoter exists upstream of this asTSS. Sequence alignment of this region in other enterobacteria shows a good conservation of a promoter with an extended -10 5’-TG box (38), suggesting that this promoter is conserved and active (Fig. 3-A).

**Figure 3:**
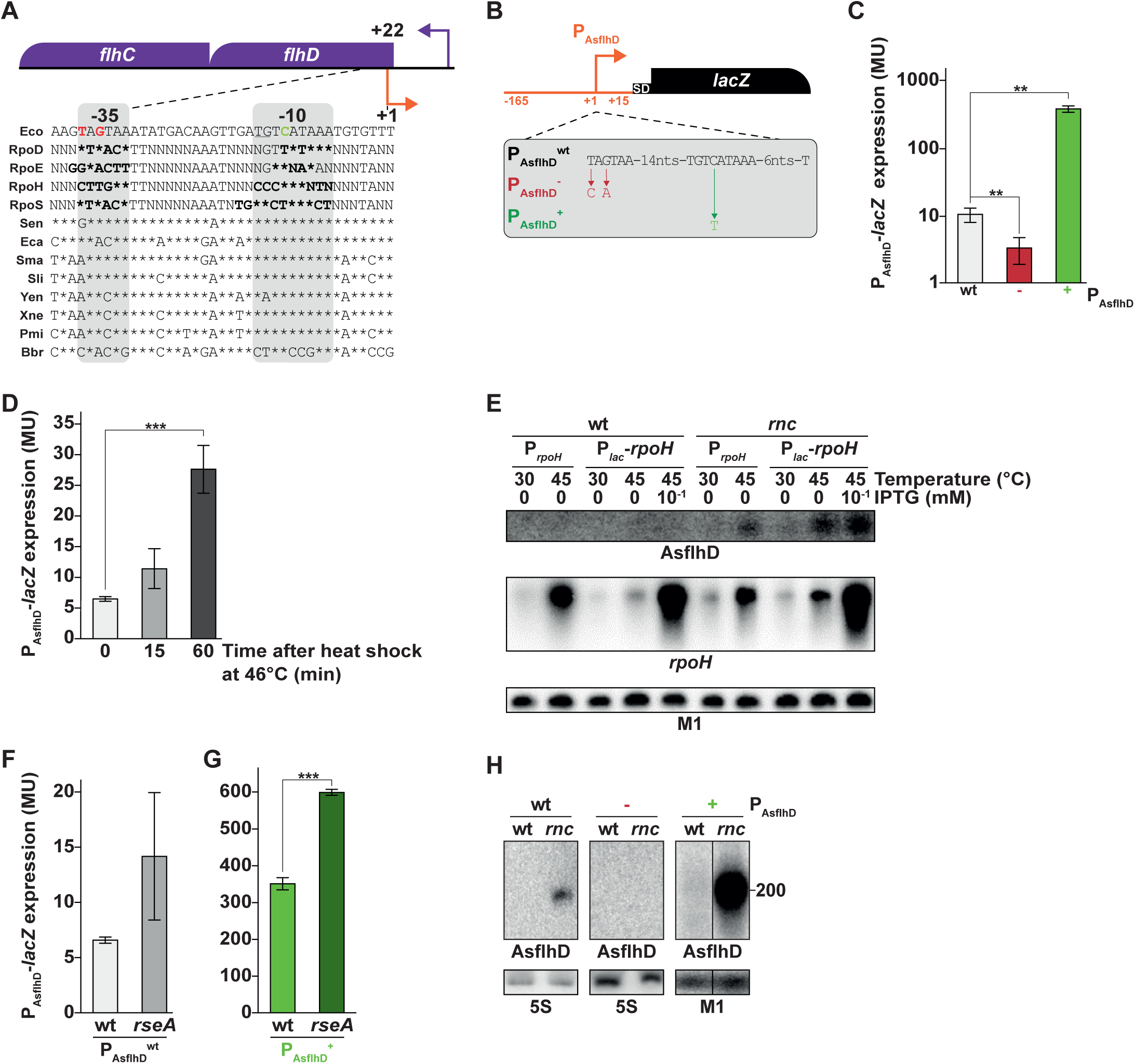
Transcriptional regulation of AsflhD. (A) Genetic structure of the *flhD* locus and alignment of the promoter sequence of AsflhD with the consensus sequences for the RpoD, RpoE, RpoH and RpoS-dependent promoters (39, 40, 77) and with 8 Eubacterial species showing between 49-92 % identity of FlhD with *E. coli* (51). The position of the promoter of AsflhD (orange bent arrow), is indicated relative to the *flhD* translation start of *flhD* (+22). Nucleotide sequences correspond to the following bacteria, Eco, *Escherichia coli* MG1655 (NC_000913.3), Sen, *Salmonella enterica typhimurium* (D43640), Eca, *Erwinia carotovora* (AF130387), Sma, *Serratia marcescens* (AF077334), Sli, *Serratia liquefaciens* (Q7M0S9), Yen, *Yersinia enterocolitica* (AF081587), Xne, *Xenorhabdus nematophilus* (AJ012828), Pmi, *Proteus mirabilis* (U96964), Bbr, *Bordetella bronchiseptica* (U17998). (B) Genetic structure of the transcriptional AsflhD-*lacZ* reporter fusion (MG2114 P_AsflhD_). Mutations in red and in green were introduced to inactivate the promoter of AsflhD (P_AsflhD_^-^ in the strain ML239) and and to increase its activity (P_AsflhD_^+^ in the strain ML218) respectively. (C) Effect of mutations in AsflhD promoter on expression of P_AsflhD_-*lacZ* fusion. (D) Expression of P_AsflhD_-*lacZ* fusion in the wt (strain (MG2114 P_AsflhD_) before (30°C t=0) and after 15 and 60 minutes of upshift (46°C). (E) DJ624, DJ624-*rnc*105 and their P*_lac_*-*rpoH* derivatives were grown at 30 °C. At mid-log phase, part of the cultures were shifted to 45°C with or without simultaneous addition of 0.1 mM IPTG. Sampling was performed 15 min later. Total RNA was analyzed by Northern blot, the membrane was probed for *rpoH*, AsflhD and M1. (F) Expression of P_AsflhD_-*lacZ* fusion in the wt strain (MG2114 P_AsflhD_) and *rseA* mutant (ML279) at 37°C. (G) Expression of P_AsflhD_-*lacZ* fusion in the strain carrying the P_AsflhD_^+^ fusion (ML218) and its *rseA* derivative strain (ML312) at 37°C. Values are means of three biological replicates for each strain, and error bars are standard deviations. Statistical significance was determined by a heteroscedastic two-tailed t test (** for p-values ≤0.01 and *** for p-values ≤0.001). (H) MG1655-B (wt), ML73 (P_AsflhD_^-^) and ML241 (P_AsflhD_^+^) and their *rnc* derivatives (respectively ML65, ML75 and ML341) were grown at 37°C until mid-log phase. Total RNA was analyzed by northern blotting. The membrane was probed successively for AsflhD and 5S or M1.

To validate the presence of a functional promoter, a P_AsflhD_-*lacZ* transcriptional fusion (P_AsflhD_^wt^) was constructed containing 165 nts before and 15 nts after the putative TSS of AsflhD (Fig 3-B). This fusion showed a relatively low level of β-galactosidase activity (Fig. 3-C). Its expression was strongly increased when the -10 motif was improved towards the RpoD consensus (P_AsflhD_^+^) while mutating the -35 to a less consensus sequence (P_AsflhD_^-^) decreased expression 2-fold (Fig. 3-C) confirming that we had identified the AsflhD promoter. It should be noted that mutations were designed to be used in the endogenous *flhD* locus, and chosen to minimally affect the coding sequence of *flhD* and to avoid introduction of rare codons. The low level of expression made us wonder if AsflhD was expressed using an alternative sigma factor. A heat-shock increased P_AsflhD_-*lacZ* expression 2-fold after 15 minutes and 5-fold after 60 minutes and also increased the level of the AsflhD RNA in the *rnc* strain (Fig. 3-DE). Comparison of the *asflhD* promoter with the consensus sequences for the two heat-shock sigma factors, σ^H^ and σ^E^ (*rpoH* and *rpoE*) shows better correlation with the σ^E^ consensus than with σ^H^ (Fig. 3-A) (39–40).

We then examined whether the P_AsflhD_ promoter is under the control of RpoE by using a strain deleted for *rseA* (anti-σ factor inhibitor of RpoE), which leads to strong induction of the RpoE regulon (41–42). Deletion of *rseA* increased 2-fold the expression of the wt P_AsflhD_-*lacZ* fusion and of the improved P_AsflhD_^+^-*lacZ* construct (Fig. 3-FG) comparable with the effect of the heat-shock at 46°C, known to induce the RpoE regulon (43). To test for an effect of *rpoH*, we introduced the P*_lac_* promoter in front of the endogenous *rpoH* gene and compared AsflhD RNA levels in wt and *rnc* mutant cells after 15 minutes of heat-shock at 45°C, in the presence or absence of IPTG. *rpoH* mRNA expressed from its own promoter or from the P*_lac_* promoter in the presence of IPTG was strongly increased by the heat-shock (Fig. 3-E). AsflhD was only detected in the *rnc* mutant and there was a correlation between the levels of *rpoH* and AsflhD transcripts at 30°C and 45°C with and without IPTG (Fig. 3-E). However, RpoH overexpression at 37°C revealed a slight repression of the transcription of AsflhD on the wt P_AsflhD_-*lacZ* fusion (Fig. S1-A) even though RNase III-stabilized AsflhD accumulated when RpoH was induced at 45°C. This would seem to rule out a direct role for RpoH in AsflhD transcription. We also tested RpoS overexpression but found it had no effect on the transcription of AsflhD (Fig. S1-B). These experiments indicate that P_AsflhD_ is functional and induced during a heat-shock due primarily to the activity of RpoE, acting directly or indirectly on the promoter of AsflhD.

To confirm the identification of the AsflhD promoter and to allow variation of the AsflhD expression, P_AsflhD_^-^ and P_AsflhD_^+^ were introduced at the endogenous *flhDC* locus and the expression of the AsflhD asRNA was examined by northern blot. The inactivation of the native promoter prevented the detection of AsflhD in the *rnc* mutant bacteria. Conversely, the mutation overexpressing AsflhD, led to the detection of a faint smear in the wt strain and to the accumulation of a high level of AsflhD in the mutant (Fig. 3-H). In summary, we have identified the AsflhD promoter and shown that the mutations in the promoter of AsflhD can be used as tools to study the function of AsflhD at the genomic locus of *flhD*.

### Characterization of the 3’-end of AsflhD

AsflhD RNA is only detected in the *rnc* strain, implying that it is very labile when RNase III is active (Fig. 3-E). Circular RT-PCR experiments (cRT-PCR) confirmed that AsflhD is indeed expressed in both the wt and *rnc* strains from the predicted asTSS promoter but that the 3′-extremities of the different AsflhD transcripts are highly heterogeneous and can extend up to 345 nts in the mutant (Fig. 4-A). Surprisingly, no 220 nts long RNA (Figs. 1-C, 3-EH) was detected in the mutant by cRT-PCR while a 149 nts long fragment was found several times exclusively in the wt strain, which might suggest that it is an intermediate in the degradation of AsflhD. It should be noted that the requirement for a ligation step during the cRT-PCR may lead to a bias towards more accessible single-stranded RNA fragments and could have excluded double-stranded RNAs from this analysis. The stable AsflhD 220 nts transcript detected in *rnc* (Fig. 5-A) should correspond to duplex RNA formation between transcripts of the convergent *flhD* and AsflhD promoters. The equivalent sense-transcript is also detected (Figs. 1-D, 5-B below). RNase III is clearly a major factor in the degradation of AsflhD. We also investigated the role of RNase E, the major endonuclease involved in mRNA turnover in *E. coli*, and of PNPase an exoribonuclease negatively controlled by RNase III. We found that the low stability of AsflhD is independent of the activity of PNPase but depends on RNase E, since a longer transcript of an approximate size of 300 nts is detected upon RNase E inactivation (Fig. 4-B). Hence, RNase III and RNase E are both involved in the rapid turnover of AsflhD but they act independently.

**Figure 4:**
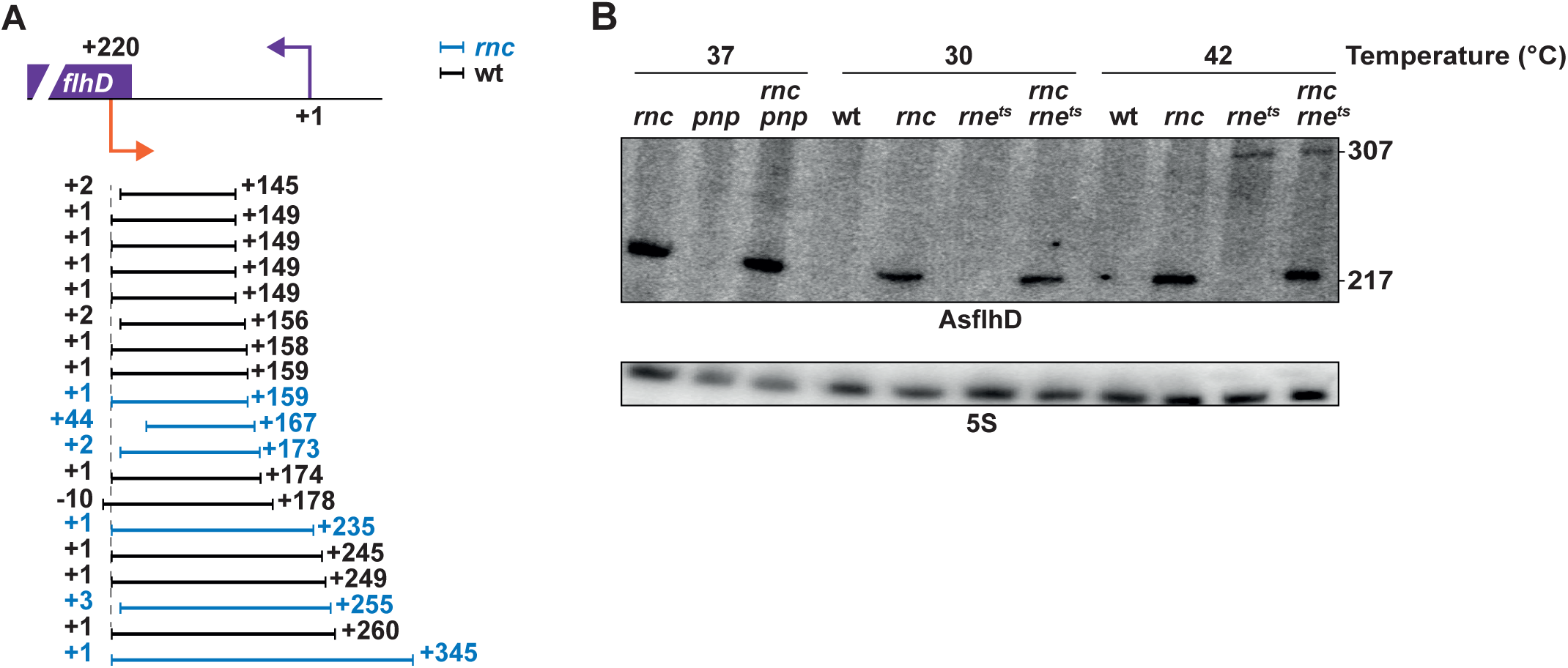
Characterization of AsflhD. (A) The sequenced reads following cRT-PCR are shown relative to the *flhD* locus. Each line represents one transcript. Transcripts were identified from both the wt strain (N3433) (in black) and from the *rnc* mutant (IBPC633) (in blue). The 5’ and 3’-end positions are indicated relative to the TSS of AsflhD. The scheme shows the localization of the P_AsflhD_ transcript relative to P*_flhD_*. (B) N3433 (wt) and its derivatives, *pnp*, *rnc*, *pnp*-*rnc*, *rne^ts^*, *rnc*-*rne^ts^* mutants (respectively N3433-*pnp*, IBPC633, IBPC633-*pnp*, N3431, IBPC637), were grown at 37°C until mid-log phase. Where indicated, cells were grown at 30°C and submitted to a heat-shock at 42°C for 15 min, in order to inactivate RNase E in the strain carrying the thermosensitive *rne*^ts^ allele. Total RNA was analyzed by northern blotting. The membrane was probed successively for AsflhD and 5S.

**Figure 5:**
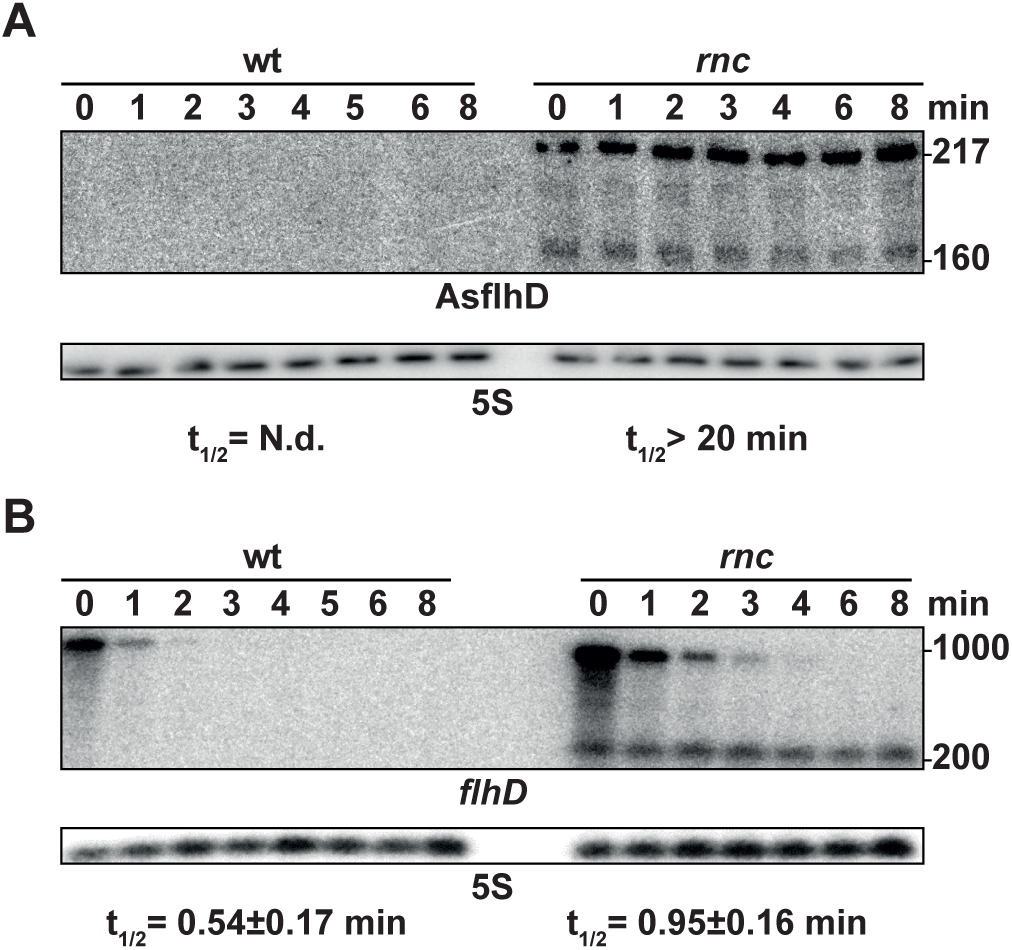
RNase III is involved in the degradation of AsflhD asRNA and flhD mRNA in vivo. (A) MG1655-B and its *rnc* derivative (ML65) were grown to mid-log phase at 37°C. At A_600_ 0.4 rifampicin (500 μg/mL) was added (t0) and sampling was performed at different times. Total RNA was extracted and subjected to northern blot analysis. The membranes were probed for AsflhD, *flhD* and 5S. The decay-rate of *flhD* mRNA was calculated as described in material and methods.

### Independent degradation of flhD and AsflhD transcripts by RNase III

We further investigated the role of RNase III in the degradation of *flhD* mRNA and AsflhD asRNA. First, we analyzed the stability of both AsflhD and *flhD* transcripts upon RNase III inactivation. In the mutant the 220 nts long AsflhD transcript and also a somewhat shorter about 160 nts long transcript were strongly stabilized, while both the amount and the stability of the long *flhDC* mRNA increased only 2-fold (Fig. 5-AB). In addition, a 220 nts long *flhD* RNA fragment was highly stabilized in the *rnc* strain. It is derived from the 5′-UTR, where the probe used in this study to detect *flhD* mRNA is located (Table S2). It presumably corresponds to a fragment of the *flhD* mRNA extending from its promoter to the AsflhD promoter located in the beginning of the *flhD* ORF (Fig. 3-A) and thus is complementary to AsflhD. The interaction of the 5’-UTR of *flhD* mRNA with AsflhD should generate an RNA duplex (Fig. 4-A) whose degradation depends on cleavage by RNase III.

We examined the interaction between AsflhD and *flhD* RNAs and their cleavage by RNase III *in vitro*. A 308 nts long *flhD* transcript corresponding to the 5’-UTR and part of the ORF of the *flhD* mRNA and a 256 nts long AsflhD asRNA were synthesized and labeled at their 5’-extremity. These two RNAs form a duplex when present in equimolar concentrations, which is completely degraded upon addition of RNase III (Fig. S2-AB). Remarkably, under the same condition, RNase III cleaves the individual RNAs independently at 2 sites on AsflhD and 4 sites on *flhD* (Fig. S2-CD). These cleavage sites are located within regions able to form secondary structure on each molecule (6, 44) (Fig. S2-C). RNase III is thus able to process both AsflhD and *flhD* RNAs *in vitro*, at specific sites but is also able to drive the complete degradation of the 5’-UTR of *flhD* when it is bound to the asRNA AsflhD. As AsflhD is never detected in the wt strain, this implies that it immediately base-pairs with *flhD* and both are degraded, so changes in AsflhD expression will directly modulate the level of *flhD* mRNA.

### AsflhD represses the expression of flhD

To investigate the function of AsflhD we determined the effect of AsflhD silencing and overexpression on *flhD* expression by following *flhD* mRNA abundance and stability using the endogenous P_AsflhD_ mutations described above. While a slight decrease (35%) of *flhD* mRNA abundance results from both silencing (P_AsflhD_^-^) and overexpression (P_AsflhD_^+^) of AsflhD, the stability of *flhD* mRNA was not significantly affected in either P_AsflhD_ mutants (Table 1).

**Table 1:**
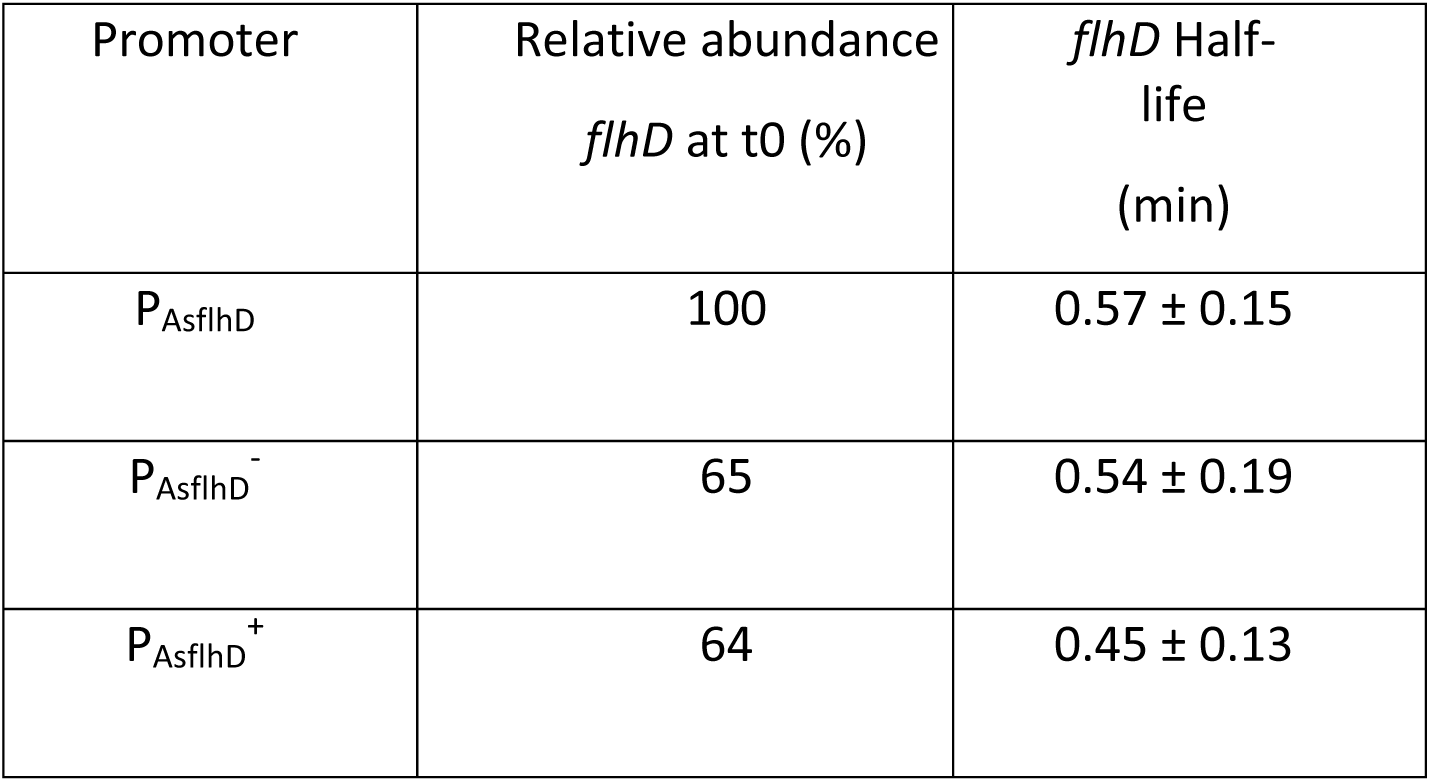
Effect of AsflhD silencing and overexpression on flhD expression and stability. MG1655-B (P_AsflhD_), ML73 (P_AsflhD_^-^) and ML241 (P_AsflhD_^+^) were grown to mid-log phase (A_600_ 0.4) at 37°C. Sampling was performed at different times after addition of rifampicin (500 μg/mL) (t0) and total RNA was subjected to northern blot analysis. The membranes were probed successively for *flhD* and M1. The decay-rate of *flhD* mRNA was calculated as described in the “quantification and statistical analysis” section of the supplementary materials and methods.

Two translational *lacZ* reporter fusions encompassing the 5’-UTR and the first 34 amino-acids of FlhD (including P_AsflhD_) were introduced at the *lacZ* chromosomal locus. The P*_flhD_*-*flhD*-*lacZ* fusion allows simultaneous monitoring of the transcriptional and translational regulation of *flhD* and the P*_tet_*-*flhD*-*lacZ,* monitors only the post-transcriptional regulation (Fig. 6-A). The mutations in the AsflhD promoter producing silencing and overexpression of AsflhD, were also introduced into both fusions. While loss of AsflhD (P_AsflhD_^-^) had no impact on the expression of FlhD, overexpression of AsflhD (P_AsflhD_^+^) resulted in decreased expression of *flhD-lacZ* expression from both the native and P*_tet_* promoters (Fig. 6-BC). Hence, overexpression of AsflhD leads to the reduction of *flhD* expression irrespective of its promoter which suggests that AsflhD is involved in the direct regulation of *flhD* mRNA and/or translational levels and not *via* an effect on the *flhD* promoter.

**Figure 6:**
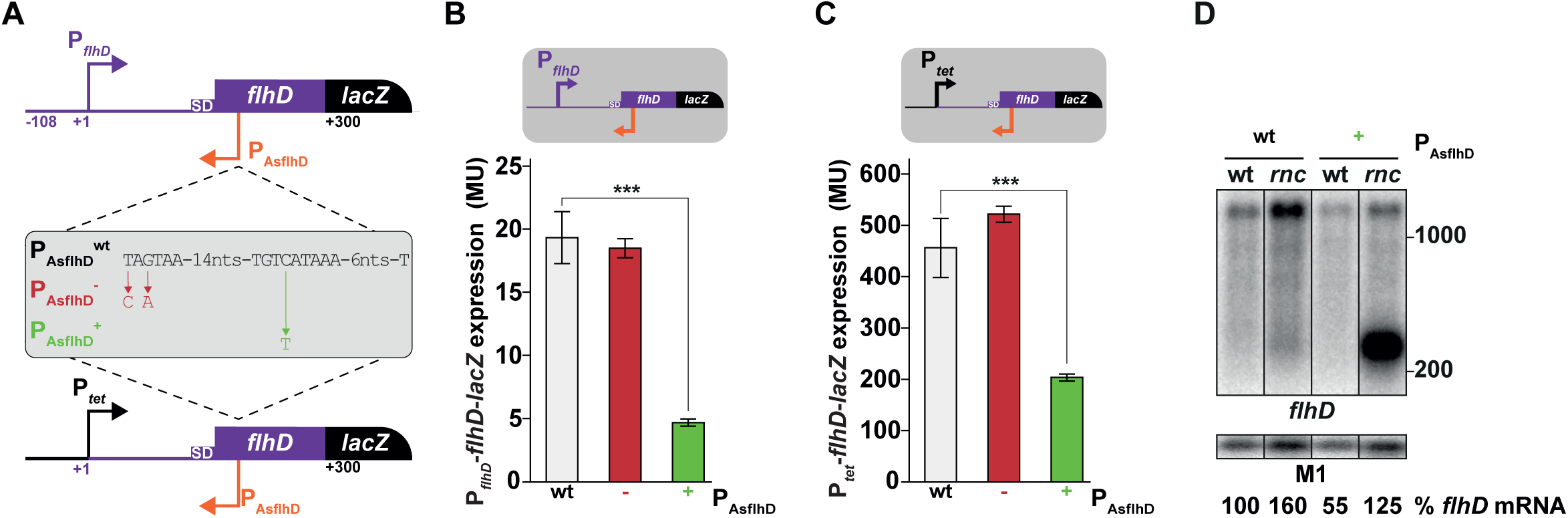
AsflhD repress the expression of flhD. (A) Genetic structures of the P*_flhD_*-*flhD*-*lacZ* (ML219) and P*_tet_*-*flhD*-*lacZ* reporter fusions (ML233) and their derivatives containing the mutations leading to either silencing (P_AsflhD_^-^, in red, ML221 and ML235 respectively) or *cis*-overexpression (P_AsflhD_^+^, in green, ML226 and ML237 respectively) of AsflhD. Expression of (B) P*_flhD_*-*flhD*-*lacZ* and (C) P*_tet_*-*flhD*-*lacZ* reporter fusions (gray bars) and their derivatives (P_AsflhD_^-^ in red and P_AsflhD_^+^ in green) are given as β-galactosidase activity. Values are means of three biological replicates for each strain, and error bars are standard deviations. Statistical significance was determined by a heteroscedastic two-tailed t test (*** for p-values ≤0.001). (D) MG1655-B (wt), ML241 (P_AsflhD_^+^) and their *rnc* mutant derivatives (ML65 and ML341 respectively) were grown to mid-log phase (A_600_ 0.4) at 37°C. Total RNA was extracted and subjected to northern blot analysis. The membrane was probed successively for *flhD* and for M1.

Finally, we determined the effect of RNase III inactivation on *flhD* mRNA in the mutant overexpressing AsflhD (P_AsflhD_^+^). Northern blot confirms that, as expected, the abundance of *flhD* mRNA increases in the *rnc* strains overexpressing or not AsflhD (Fig. 6-D). Remarkably, the 220 nts long fragment observed upon RNase III inactivation (Figs. 4-B, 5-B) strongly accumulates when AsflhD is overexpressed (Fig. 6-D). Hence this suggests that the increase in strength of the AsflhD promoter drives the accumulation of the small fragment corresponding to the 5’-UTR of *flhD* mRNA, presumably as a duplex with AsflhD and that both are rapidly degraded by RNase III. This short *flhD* fragment could either be generated by processing of longer *flhD* mRNA or correspond to premature transcriptional termination of *flhD* mRNA. In summary, we show that AsflhD is involved in the repression of the expression of *flhD* at the post-transcriptional level.

### Mutual repression of transcriptional elongation by AsflhD and flhD in vitro

To determine whether AsflhD can repress the transcription of *flhD*, we performed *in vitro* transcription experiments using a DNA template corresponding to the *flhD* gene from 76 nts before to 388 nts after the transcription start site of *flhD*, which allows the transcription of a 388 nts *flhD* RNA and of a 335 nts AsflhD RNA (Fig. 7-A). We compared the abundance of both transcripts synthesized from this latter DNA fragment to those generated from templates carrying the promoter mutations leading to either silencing (P_AsflhD_^-^) or overexpression (P_AsflhD_^+^) of AsflhD. *In vitro* transcription assays were performed in a single round of elongation in the presence of heparin and with RNA polymerase (RNAP) pre-bound to templates in the absence of RNA, hence observed effects are restricted to the elongation step and should be independent of the initiation of transcription. The results correlate with *in vivo* data even though the amplitude of the effects is different. Figure 7 shows that expression of AsflhD is strongly impaired (10-fold) on the template carrying the silencing (P_AsflhD_^-^) mutation and increased (2.5-fold) from the template carrying the overexpression (P_AsflhD_^+^) mutation of AsflhD (Fig. 7-B orange bars). Also, as *in vivo* (Fig. 6-B), AsflhD silencing does not affect the level of the *flhD* RNA, while its overexpression in *cis* results in a decrease of the transcription of the *flhD* RNA (40%) (Fig. 7B purple bars). cAMP/CAP is known to activate the transcription of *flhD* by binding to a sequence located 72 nts upstream from the TSS of *flhD* (45). As expected, its addition increased the transcription of *flhD*, which was still reduced by AsflhD overexpression (Fig. 7-B left). The role of AsflhD in the repression of *flhD* transcription elongation, was further confirmed by using a template where the P*_tet_* promoter replaced the P*_flhD_* promoter (Fig. S3-A) producing the same 388 nts *flhD* RNA but a shorter (260 nts) AsflhD RNA. Similar results were observed from the templates carrying the P_AsflhD_ silencing and overexpression mutations (Fig. S3-B). We conclude that overexpression of AsflhD *in cis* leads to the repression of transcription elongation of *flhD,* which seems to be independent of the transcription level and of the promoter expressing *flhD*.

**Figure 7:**
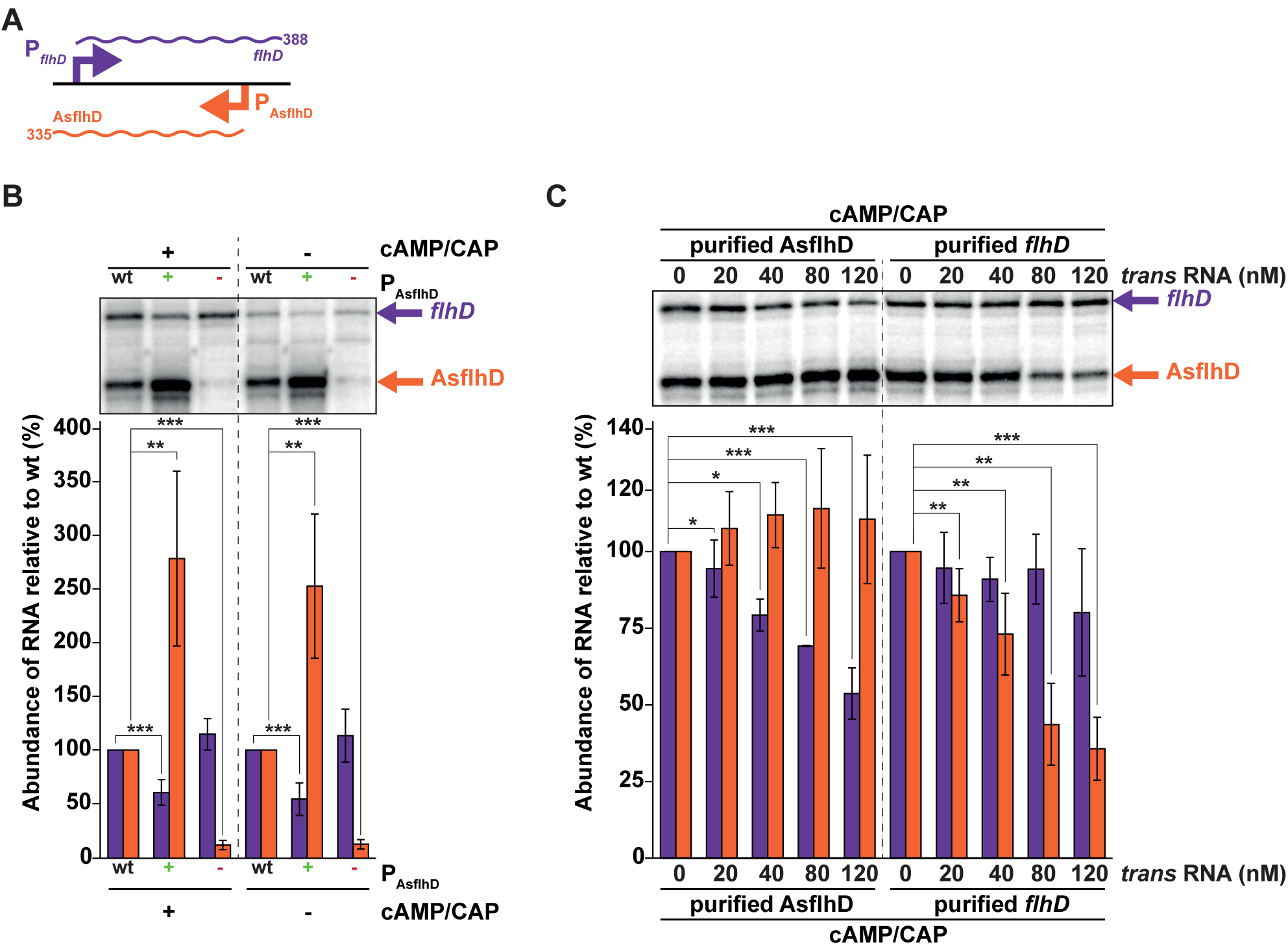
AsflhD is involved in the transcriptional attenuation of flhD. (A) Schematic representation of the template used for the *in vitro* transcription assay carrying the P*_flhD_* promoter driving the expression of a 388 nts transcript (purple) and the P_AsflhD_ promoter driving the expression of a 335 nts transcript (orange). The linear DNA template was constructed using the oligonucleotides LM191 and LM9 (table S1) and corresponds to -76 to +388 of the *flhD* transcript relative to its TSS, with a 40 nts extension carrying the *rrnB*T2 terminator (fragment length 504 bp). This fragment carries the native *flhD* promoter (-10 and -35 sites) and includes the cAMP/CAP site at -72 compared to the *flhD* TSS, at its upstream extremity. *In vitro* transcription assays were performed as described in the Supplementary material and method (B) with or without addition of 100 nM CAP and 0.2 mM cAMP for 15 min at 37°C before addition of RNA polymerase, (C) with 100 nM CAP and 0.2 mM cAMP and the addition of *in vitro* purified AsflhD or *flhD* transcripts to the reaction at the indicated concentrations. Samples were analyzed on sequencing gels. Relative intensity of the indicated bands (*flhD* in purple and AsflhD in orange) were analyzed. Values are means of 6 (B) or 3 (C) replicates and error bars are standard deviations. Statistical significance was determined by a heteroscedastic two-tailed t test (* for p-values ≤0.05, ** for p-values ≤0.01 and *** for p-values ≤0.001).

We then determined the effect of purified AsflhD or *flhD* RNA addition on the transcription of both *flhD* and AsflhD using the same linear DNA templates. Figure 7-C shows that addition of increasing amount of AsflhD led to a linear decrease of *flhD* while not affecting the accumulation of AsflhD. The reciprocal assay by adding increasing concentrations of purified *flhD* RNA decreased linearly the amount of AsflhD synthesized, while the amount of *flhD* was not affected. We performed the same assay with the shorter template carrying the P*_tet_* promoter and observed similar results (Fig. S3-C). In summary, AsflhD represses the transcription elongation of *flhD* both in *cis* and in *trans*, independently of the promoter and its expression level. Thus, we propose that AsflhD asRNA and *flhD* mRNA are involved in their mutual transcriptional attenuation in which the interaction of one molecule with the other leads to a reduction in transcription *via* an alteration of transcription elongation.

### AsflhD represses the transcription of flhD in trans in vivo

To confirm *in vivo* the ability of AsflhD to repress the transcription of *flhD* in *trans*, AsflhD was overexpressed from a plasmid, under the control of a P*_tac_* promoter inducible by IPTG. The short 242 nts long AsflhD is transcribed from the +1 to the +220 nt relative to the TSS of AsflhD with a *rrnB*T2 terminator to enable its stabilization. In the *rnc* mutant, this transcript is further processed with the appearance of the characteristic 160 nts intermediate (Figs. 8-A, 1-D). It is noteworthy that this smaller fragment is slightly longer than in the endogenous AsflhD suggesting that it may correspond to a 3’-fragment of AsflhD. Overexpression of AsflhD upon addition of IPTG decreases the abundance of *flhD* mRNA both in the wt strain (30%) and in the mutant (20%) (Fig. 8-B), in agreement with our *in vitro* data (Figs. 7-C, S3-C). Thus, consistent with previous experiments, *trans* overexpression of AsflhD *in vivo* reduces the abundance of *flhD* mRNA but it is less effective than AsflhD expressed from the *flhD* locus *in cis* and this effect is mostly independent of RNase III activity.

**Figure 8:**
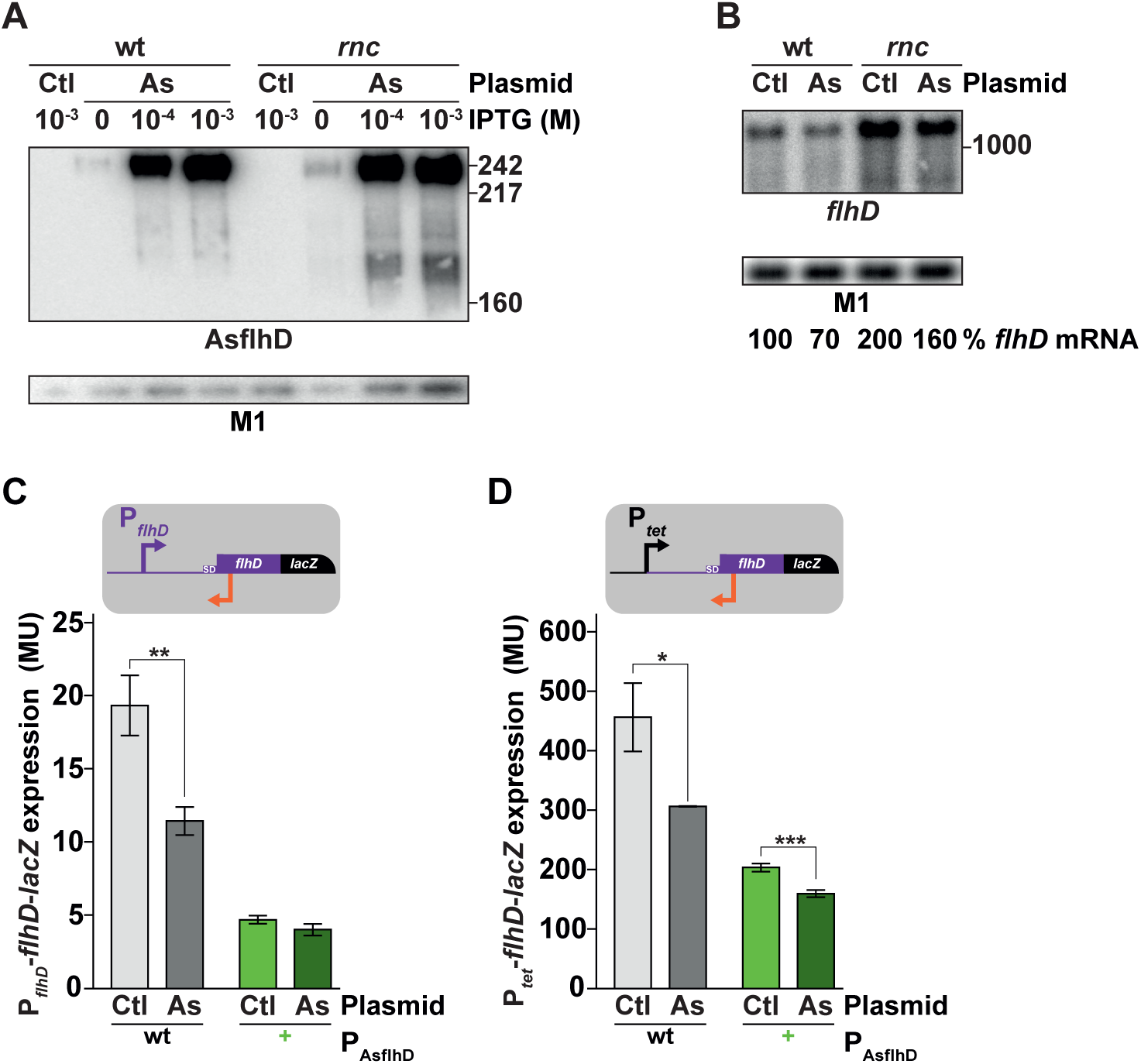
AsflhD repress the expression of flhD in trans. (A) MG1655-B (wt) and its *rnc* derivative (ML65) containing the control pCA24N (Ctl) or the pCA24N AsflhD (As) plasmids were grown to mid-log phase (A_600_ 0.4) at 37°C in the presence of the indicated concentration of IPTG. Total RNA was extracted and subjected to northern blot analysis. The membrane was probed for AsflhD and M1 RNA. (B) MG1655-B (wt) containing the control pCA24N (Ctl) or the pCA24N AsflhD (As) plasmids were grown to mid-log phase (A_600_ 0.4) at 37°C in the presence of 10^-4^ M IPTG. Total RNA was extracted and subjected to northern blot analysis. The membrane was probed for *flhD* using a probe corresponding to the beginning of the *flhD* ORF and M1 RNA. This *flhD* probe was used to detect *flhD* from the chromosomal locus and to avoid detection of *flhD* RNA transcribed from a plasmid promoter. Expression of (C) P*_flhD_*-*flhD*-*lacZ* (ML219) and (D) P*_tet_*-*flhD*-*lacZ* (ML233) reporter fusions (gray bars) and their P_AsflhD_^+^ derivatives (in green, ML226 and ML237 respectively) containing the pCA24N control (Ctl, in dark grey) or the pCA24N AsflhD (As, in dark green) plasmid was determined in the presence of 10^-4^ M of IPTG. Values are means of three biological replicates and error bars are standard deviations. Statistical significance was determined by a heteroscedastic two-tailed t test (* for p-values ≤0.05, ** for p-values ≤0.01 and *** for p-values ≤0.001).

To further confirm that AsflhD can impact *flhD* expression *in vivo*, we measured the expression of the *flhD*-*lacZ* reporter fusions when AsflhD is overexpressed from the plasmid. The overexpression induces about a 30-40% decrease in the expression of *flhD* both from the P*_flhD_*-*flhD*-*lacZ* and the P*_tet_*-*flhD*-*lacZ* fusions (Fig. 8-CD, dark grey). However, overexpression of AsflhD from the plasmid in the strain where expression of AsflhD is already upregulated, by the presence of P_AsflhD_^+^ mutation in *cis,* has no or little additive effect on the final repression (Fig. 8-CD, dark green). It should be emphasized that these effects are independent of the *flhD* promoter (native P*_flhD_* or P*_tet_*). In summary, we demonstrate that AsflhD RNA is involved in the transcriptional attenuation of *flhD* both in *cis* and in *trans*, and that it does not involve the native P*_flhD_* promoter.

### AsflhD controls the motility cascade

The *flhDC* operon encodes the FlhD_4_C_2_ transcriptional master regulator of the swimming motility, so we next investigated the effect of AsflhD overexpression on the expression of key factors belonging to the motility cascade. This cascade of gene activation is divided into three classes of genes (46). The *flhDC* operon encodes the only Class I protein, FlhD_4_C_2_, which is required for expression of class II genes, which in turn control class III genes. We tested the effect of AsflhD on representative Class II and Class III genes. The selected class II genes are *fliA* that encodes FliA, the sigma factor for class III motility genes, and *flgB* that encodes FlgB, the main component of the flagella rod. The *fliC* gene is a class III gene, located upstream from the *fliA* gene, and it encodes the main component of flagella, FliC. The amounts of *fliA*, *flgB* and *fliC* mRNAs are all very strongly reduced upon overexpression of AsflhD from its endogenous locus in the mid-log phase (Fig. 9-ABC). As the reduction is considerably stronger than the effect on *flhD* itself, this implies that the effect of AsflhD overexpression is amplified compared to that on *flhD,* similarly to what was previously reported for transcriptional regulators of *flhD* expression (47–48). Furthermore, a bioinformatics search (TargetRNA2 (49)) for possible direct *trans* targets of AsflhD found no candidates amongst genes from the motility cascade. In addition, the abundance of *flgB* and *fliC* mRNAs in late-log phase slightly increases relatively to mid-log phase (Fig. S4-AB) therefore this is consistent with the hypothesis that *cis*-overexpression of AsflhD limits the level but also delays the timing for the induction of the motility cascade.

**Figure 9:**
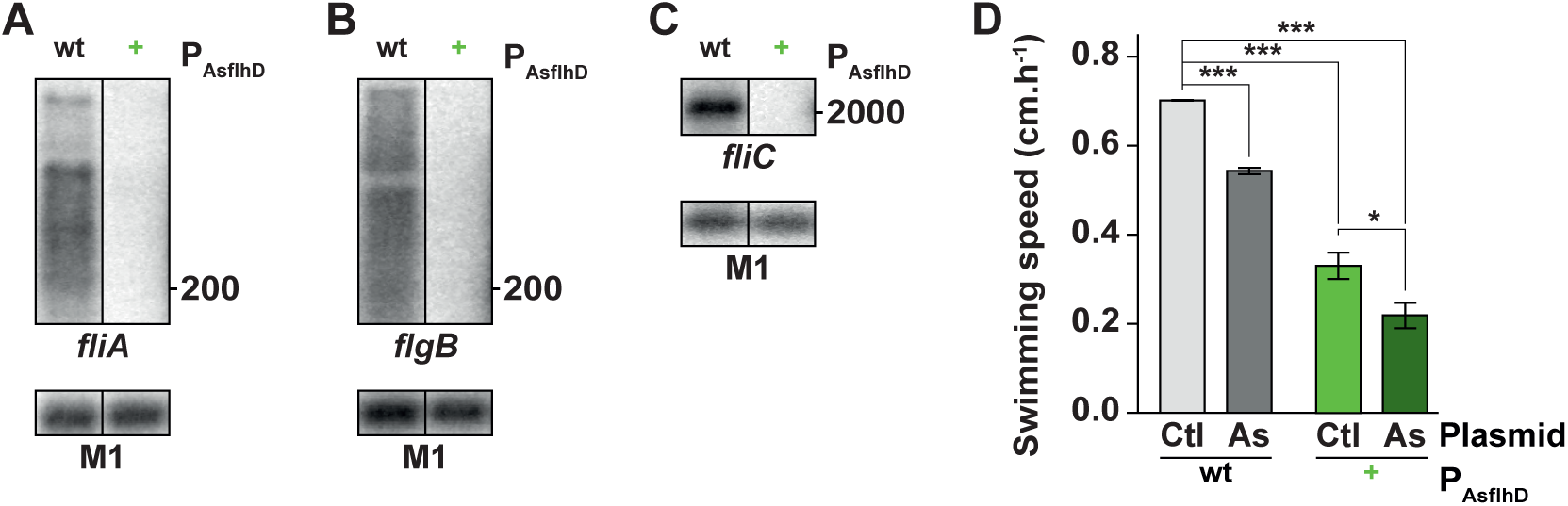
AsflhD overexpression represses the cascade of motility. MG1655-B (wt) and ML241 (P_AsflhD_^+^) were grown at 37°C until mid-log phase. Total RNA was analyzed by northern blotting. The membranes were probed successively for (A) *fliA* and M1, for (B) *flgB* and M1 and for (C) *fliC* and M1. (D) Swimming motility speed is reduced upon overexpression in *cis* (P_AsflhD_+ green bars) and in *trans* (pCA24N AsflhD dark grey and dark green bars) of AsflhD as observed on motility plates. Swimming speed (cm.h^-1^) was calculated (see quantification and statistical analysis section of the supplementary material and method) on three biological replicates in the wt strain (MG1655-B) and in the P_AsflhD_^+^ strain (ML241) carrying the pCA24N control (Ctl) or the pCA24N AsflhD (As) plasmid. Values are means of three biological replicates and error bars are standard deviations. Statistical significance was determined by a heteroscedastic two-tailed t test (* for p-values ≤0.05, ** for p-values ≤0.01 and *** for p-values ≤0.001).

Finally, we analyzed the effect of AsflhD on the motility of bacteria on low-agar plates and observed that overexpression of AsflhD from both the *flhD* locus (P_AsflhD_^+^) or from the plasmid decreases the swimming speed on plates (50% and 30% respectively) while the combined *cis* and *trans* overexpression of AsflhD induces a stronger reduction of the swimming speed (70%) (Fig. 9-D). These results enforce our hypothesis that the observed strong reduction of the motility cascade effectors (*fliA*, *flgB* and *fliC* genes) in mid-log phase reflects a delayed induction rather than actually inhibiting expression.

In summary, both *cis* and *trans* overexpression of AsflhD reduces transcription of *flhD*, which in turn leads to repression of the whole cascade of motility and a reduction in swimming speed. Of note, *cis-*expressed AsflhD appears to be more effective than the *trans-*expressed to control *flhD* expression (Fig. 8-CD) and bacterial motility (Fig. 9-D). We hypothesize that the overexpression of AsflhD delays the induction of *flhD* to maintain the timing of the motility program.

## Discussion

Regulatory RNA molecules are often part of complex genetic networks in bacteria. They correspond to a heterogeneous class of molecules that differ in gene organization, size and function. Our goal was to detect, identify and investigate the function of some antisense transcripts in *E. coli*. We compared the transcriptomes of an *rnc* mutant to that of a wt strain and selected candidate asRNAs. We validated the presence of discrete transcripts, only detected in the *rnc* mutant, which were complementary to *crp*, *ompR*, *phoP* and *flhD* genes. We identified the promoters encoding AsphoP and AsflhD, both producing transcripts convergently expressed towards the promoters of their target genes *phoP* and *flhD.* We show that RNase III is involved in the decay of both AsphoP and *phoP.* AsflhD is highly unstable and it is degraded independently by RNase E and RNase III. The promoter of AsflhD is induced by a heat-shock and is partially dependent on RpoE. We reveal that AsflhD is involved in transcriptional attenuation of *flhD* by being able to repress *flhD* transcription both *in vitro* and *in vivo* when overexpressed in *cis* or in *trans* (Figs. 7-8). Remarkably, *in vitro, flhD* RNA in *trans* could also drive the transcriptional attenuation of AsflhD asRNA (Fig. 7), suggesting that the interaction of the two RNAs can perturb the transcription elongation of the other transcript *in vivo*. The relatively small effect of AsflhD on *flhD* expression has important consequences for the FlhD_4_C_2_ regulon since, the *cis* overexpression of AsflhD lead to decreased expression of three representative genes of the motility cascade; *fliA, flgB* and *fliC*, and reduced the swimming speed (Fig. 9). In conclusion, this work demonstrates that AsflhD is an additional player acting in the already complex regulatory process controlling *flhDC*. Our view of how AsflhD, by co-transcriptionally fine-tuning the expression of *flhD,* reinforces the control of the motility cascade, is shown in Figure 10.

**Figure 10:**
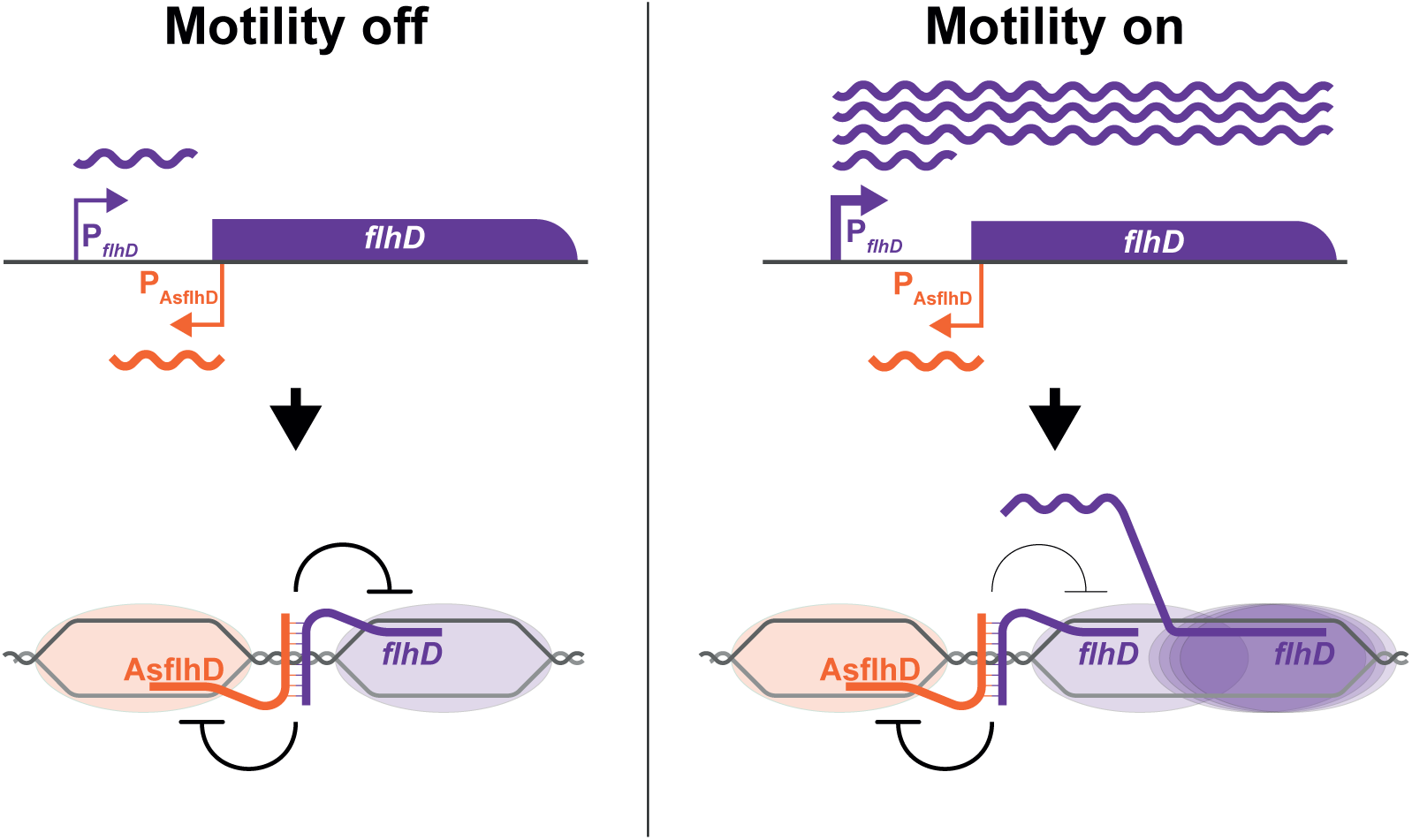
Schematic representation of the regulatory function of AsflhD. When the motility cascade is off, low expression from the P*_flhD_* promoter (purple bent arrow) is not sufficient to maintain the expression of *flhDC*. The binding of AsflhD (orange) to *flhD* (purple) RNA may occur co-transcriptionally and perturb the elongation of both molecules reinforcing the low level expression of *flhD*. Upon encountering conditions where motility is required, the strong induction of the P*_flhD_* promoter changes the balance between the sense and the asRNA allowing *flhDC* expression and activation of the motility cascade but where changes in AsflhD expression (e.g. in response to heat) could modulate *flhD* expression. DNA is represented by grey lines and RNAP as orange and purple oval shapes.

### Conservation of the promoters of AsflhD and AsphoP

asRNAs are poorly conserved among bacteria (50), but if this can be expected for asRNAs from intergenic regions with low sequence constraints, nucleotide changes within the coding region of the target risk to upset the function of the ORF and could be counterselected. AsflhD corresponds almost entirely to the 5’-UTR of *flhD* but with the promoter located in the ORF, which is fairly well conserved in enterobacteria (Fig. 3-A). The presence of multiple critical amino-acids along the protein, whose modification leads to decreased motility, could explain the conservation of the amino sequence of FlhD among Gram-negative bacteria (51). Thus, the conservation of the promoter of AsflhD could be the result of direct selection for FlhD activity or for the regulatory function of AsflhD controlling the expression of FlhD in these bacteria. In spite of the close similarity between the AsflhD promoter regions of *Salmonella enterica* sv Typhimurium and that of *E. coli* (Fig. 3-A), it would be interesting to examine whether AsflhD is also expressed and controls *flhD* levels in *S. enterica,* since it was reported that *E. coli flhDC* operon is expressed in *S. enterica* but at significantly lower levels (52).

The promoter of AsflhD shows low activity *in vivo* despite a -10 consensus with an extended -10 5′-TG-3’ element and a -35 element which is functional since the P_AsflhD_^-^ mutation decreased the promoter activity. In the case of AsphoP, the promoter located in the central region of the *phoP* ORF is not conserved in other enterobacteria. In *E. coli*, the two G to A changes in the -10 sequence of the promoter of AsphoP, compared to other bacteria make a relatively good consensus -10 (CATAAT) which can account for the high level of expression of AsphoP (Fig. 2-E) compared to AsflhD (Fig. 3-C). The lack of conservation in other bacteria suggests that, on the contrary to AsflhD, any function of AsphoP may be unique to *E. coli* where it was most likely acquired.

### Role of RNase III in the degradation of asRNAs-mRNAs

We demonstrate that RNase III initiates the rapid degradation of AsphoP and AsflhD RNAs, at the same time it is also involved in the *flhD* degradation. The formation of an intermolecular RNA duplex can trigger the RNase III-mediated degradation of a mRNA as in the case of the *puf* mRNA upon binding to asPcrL in *Rhodobacter sphaeroides* or a sRNA upon binding to its target, as in the case of the sRNA RyhB upon binding to the *sodB* mRNA (53). However, RNase III cleavage may generate a shorter but more stable mRNA and so can have a positive role on gene expression (19, 54). In the case of AsphoP the *rnc* mutation reduces promoter activity (Fig. 2-E) but greatly increases AsphoP stability (Fig. 2-A). In the case of AsflhD, RNase III allows the recycling of the products of transcriptional attenuation between *flhD* and AsflhD (Fig. 7-A).

### Mechanism of regulation by AsflhD

*Cis*-encoded regulatory elements and asRNAs have the advantage of their close location facilitating their access to their target. Binding of AsflhD to the 5’-UTR of *flhD* mRNA can have multiple consequences on the interaction of other *trans*-acting regulators, known to affect *flhD* expression. For example, after removal of the 5’-triphosphate, CsrA binds two regions of the *flhD* mRNA and protects it by competing with RNase E (6, 55). Closer to the translational start, the binding of the McaS sRNA is required to expose the ribosome binding site and activate translation. On the contrary, binding of the sRNAs OxyS, ArcZ, OmrA and OmrB represses translation (5, 56). AsflhD binding can interact at any of these sites along the *flhD* mRNA to compete with the RNase E-mediated degradation pathway and/or the sRNA control of translation. Our results clearly demonstrated that AsflhD represses the transcription of *flhD* but without excluding that AsflhD may also be involved in the control of the translation of *flhD* mRNA. However, it is difficult to assess whether AsflhD may have a direct effect on translation or an indirect effect *via* the competition with the other positive and negative post-transcriptional regulators of translation of *flhD*.

AsflhD and *flhD* mutually repress their transcription elongation. AsflhD interacting with the 5’-UTR of *flhD* mRNA, could either provoke the termination of transcription by transcriptional interference or by transcriptional attenuation (57), or drive the processing of *flhD* mRNA (*e.g., via* RNase III) or inhibit translation. Diverse mechanisms of transcriptional attenuation have been described which are dependent on regulatory RNAs. For example, sRNA binding drives the premature Rho-dependent transcription termination of the *rpoS* mRNA (58). The asRNA RNAβ in *Vibrio anguillarum* promotes transcription termination on the *fatDCBA* polycistronic mRNA before RNA polymerase reaches the end of the *fatDCBAangRT* operon. Remarkably, the region where termination occurs does not contain any canonical terminator motif (59). The asRNA RnaG was also shown to stabilize a terminator structure upon binding to the *icsA* mRNA in *Shigella flexneri* (16). Similarly, the asRNA RNAIII binds to the leader region of the *repR* mRNA and favors the formation of a terminator structure in *Bacillus subtilis* (60). The asRNA anti-Q in *Enterococcus faecalis* is responsible for both transcriptional interference due to RNAPs collisions and attenuation by an uncharacterized mechanism (61). Our experiments do not detect the accumulation of a shorter transcript *in vitro* upon addition of one or the other of the transcripts suggesting that binding of AsflhD to *flhD* does not stabilize a terminator structure but could rather modify the stability of the elongating RNAP leading to heterogenous 3’-termini as observed for AsflhD *in vivo* by cRT-PCR (Fig. 4-A).

### Physiology of AsflhD

Motility depends on the growth-rate due to the regulation of *flhD* (62). The expression of *flhD* peaks at the end of the exponential phase then decreases and is stabilized to an intermediate level during the stationary phase in *E. coli* (63). The temporal control of the motility cascade is maintained *via*, among other mechanisms, the expression of anti-σ factor, FlgM, at the same time as FliA, in order to allow the expression of class II genes without inducing class III genes expression prematurely. When the basal part of the flagellum is assembled, FlgM is exported from the cytoplasm and FliA induces the expression of class III genes.

We show here that the P_AsflhD_ promoter is induced during a heat-shock likely *via* RpoE. Remarkably, the swimming motility behavior is down-regulated during a heat-shock. This was proposed to be due to both a lowered level of FlhD and the inefficient export of FlgM (64–65). Our results show that the up-regulation of AsflhD during the heat-shock reducing *flhD* expression could also contribute to the decrease in swimming motility. Thus, as well as providing a fine-tuning mechanism to coordinate the expression of *flhD*, a function of AsflhD could be help to maintain the motility cascade off in conditions where motility would be detrimental and/or too costly.

### Outlook

Our results demonstrate that the asRNA AsflhD is involved in a mutual transcriptional attenuation mechanism with its target *flhD* mRNA. Regulatory RNAs are far from being fully understood in bacteria and new mechanisms of action are likely to be discovered. Development of global approaches able to capture RNA-RNA and RNA-protein interaction (66–67) as well as prokaryotic single cell RNA-seq (68) are likely to pave the way for the elucidation of the role of the widely distributed asRNAs which were, until recently mostly considered as pervasive transcriptional noise but for which the study of individual cells and molecules could be critical for the understanding of their function.

## Materials and methods

### Bacterial strains and culture conditions

Strains and plasmids used in this work are listed in table S1. Constructions and mutations were made by using primers given in table S2 and are described in Supplementary Materials and methods. Strains were grown in LB Miller medium at 37°C, or at 30°C and shifted to 42°C, 45°C or 46°C for the heat-shock experiments. Appropriate antibiotics were added when required. IPTG was used at the indicated concentrations for induction of AsflhD from the pCA24N AsflhD plasmid and arabinose for RpoS from the pBAD18 plasmid.

### Northern blotting and RNA-seq analysis

Total RNA was prepared from bacteria grown to the A_600_ 0.4 using the hot-phenol procedure (69). Five μg of total RNA were electrophoresed either on 1% agarose, 1xTBE or 6% polyacrylamide gels (19/1), 7M urea, 1xTBE for analysis by northern blotting (70–71) along with RiboRuler High-Range marker (ThermoFisher) or radio-labeled Msp1-digested pBR322 (NEB). Membranes were hybridized with complementary RNA probes. Templates for the synthesis of the RNA probes were obtained by PCR amplification using the pair of “m” and “T7” oligonucleotides (Table S1). Probes were synthesized by T7 RNAP with [α-^32^P]-UTP yielding uniformly labeled RNAs (72). Membranes were also probed with M1 or 5S as loading control by using 5’-end labeled primers (Table S2). An RNA-seq analysis was performed to compare the transcriptomes of the wild-type (N3433) and the RNase III deficient strain (IBPC633). Sample preparation for RNA-seq, 5′-RNA tagging and RNA-seq analysis were performed as in (25). Data have been deposited in the ArrayExpress database at EMBL-EBI under accession number E-MTAB-9507 (M. Lejars, L.Kuhn, A. Maes, P. Hammann, E. Hajnsdorf manuscript in preparation).

### β-galactosidase assays

Cultures were initiated at A_600_ 0.05 and sampled at A_600_ 0.4 or in the case of the results presented in figure S2 also at A_600_ 1.2. Samples (200 µL) were lysed in 800 µL PBS buffer with 10 µL 0.1% SDS and 20 µL chloroform. β-galactosidase activity was assayed as described (73), results are the mean of at least three biological replicates.

### Circular RT-PCR

Circular RT-PCR was performed with total RNA extracted from N3433 and IBPC633 treated with 5′-polyphosphatase. After circularization with T4 RNA ligase, mflhD2 was used to prime reverse transcription and mflhD6 and masflhD10 to generate PCR products (Table S2), which were cloned (74).

### RNA band-shift assay and in vitro processing by RNase III

DNA templates carrying a T7 promoter sequence were generated by PCR using the Term and T7 oligonucleotides (Table S2). They allow the transcription of the first 308 nts *flhD* and of the first 256 nts of AsflhD. RNAs were synthesized by T7 RNAP with [α-^32^P]-UTP as a tracer and were gel purified. Transcripts 5′-end-labelling, hybridization, RNase III digestion and sample analysis were described in (74–76). Briefly, radioactive AsflhD was incubated with increasing concentrations of *flhD* mRNA under two conditions referred to as “native” (incubation in TMN buffer (20 mM Tris acetate, pH 7.5, 10 mM magnesium acetate, 100 mM sodium acetate for 5 min at 37◦C) and “full RNA duplex” (initial denaturation at 90°C for 2 min, then incubation in 1xTE at 37°C for 30 min). The complexes were loaded on native polyacrylamide gels to control for hybridization efficiency or submitted to *in vitro* processing by RNase III of *E. coli*. RNase III digestion of free 5’-radiolabeled AsflhD, *flhD* or complexed AsflhD with *flhD* was performed at 37°C in TMN buffer containing 1 µg tRNA for 15 min with RNase III (Epicentre). Samples were loaded on denaturing polyacrylamide gels together with an RNA alkaline ladder as in (75).

### In vitro transcription assay

Single-round *in vitro* transcription experiments were carried out on linear templates as described in Supplementary materials and methods.

### Motility assay

Stationary phase bacterial cultures (MG1655-B, ML241 (P_AsflhD_^+^) carrying the pCA24N control (Ctl) or the pCA24N AsflhD (As) plasmid) were inoculated (2 µL) on soft-agar (0.2 g/L) SOB motility plates (containing 10^-4^ M IPTG and 2.4 g/L MgSO_4_) at 37°C and pictures were taken using a Gel Doc (Biorad) imager at the beginning and the end of the linear swimming motility period (from 5 to 8 hours). Swimming speed was then calculated as a function of time by comparing motility diameters.

*Image treatment, quantifications and statistical analysis* are given in Supplementary Materials and Methods

## Abbreviations

asRNA: Antisense RNA
RBP: RNA-binding protein
UTR: untranslated region
sRNA: small RNA
ORF: open reading frame
nt: nucleotide
cRT-PCR: circular RT-PCR
RNAP: RNA polymerase

## Acknowledgements

We thank C. Beloin and J-M Ghigo for providing strains and A. Kolb for the kind gifts of purified RNAP core, σ^70^ and CAP. This work was supported by the Centre National de la Recherche Scientifique (UMR8261), Université de Paris, Agence Nationale de la Recherche (asSUPYCO, ANR-12-BSV6-0007-03 to E.H.), (RIBECO ANR-18-CE43-0010 to E. H.) and the “Initiative d’Excellence” program from the French State (Grant “DYNAMO,” ANR-11-LABX-0011).

## Supplementary legends

*Figure S1: The transcription of AsflhD is not activated by RpoH and RpoS*

(A) Expression of AsflhD-*lacZ* fusion in the wt strain (MG2114 P_AsflhD_) and its P*_lac_*-*rpoH* derived strain (ML310) in the presence of 10^-3^ M IPTG at 37°C was measured at OD_600_ 1.2. (B) Expression of AsflhD-*lacZ* fusion in the wt strain (MG2114 P_AsflhD_) carrying the pBAD18 control (Ctl) or pBAD18 RpoS (*rpoS*) in the presence of 0.1% arabinose at 37°C was measured at OD_600_ 1.2. Values are means of three biological replicates and error bars are standard deviations. Statistical significance was determined by a heteroscedastic two-tailed t test (** for p-values ≤0.01).

*Figure S2: In vitro cleavage of AsflhD and flhD RNAs by RNase III*

5′-radiolabeled AsflhD RNA (308 nts) was incubated with increasing concentrations of *flhD* mRNA (256 nts) under conditions referred as Native and Full RNA duplex conditions (Materials and Methods). Native AsflhD-*flhD* complexes were formed at 37°C for 5 min in TMN buffer, and full duplexes were obtained after a denaturation-annealing treatment in TE Buffer (2 min 90°C, 30 min 37°C before loading on native polyacrylamide gels to control for hybridization efficiency (A) or (B) *in vitro* processing by RNase III. RNase III digestion of free or complexed AsflhD in native conditions was performed at 37°C in TMN buffer containing 1µg tRNA with increasing concentration of RNase III per sample. Samples were analyzed on 8% polyacrylamide-urea gels. 5′-radiolabeled *flhD* (C) and AsflhD (D) were cleaved by RNase III (1 unit) *in vitro* at 4 and 2 main sites respectively (represented by a numbered red arrow). Mapping of the main *in vitro* cleavage sites of RNase III on *flhD* and AsflhD are indicated on their predicted secondary structures with the position of the main RNase III cleavage sites indicated relative to the TSS (according to (6) and Vienna RNA websuite (44)). The localization of RNase III cleavage site (black arrows) was performed by comparing the cleavage fragment relative to an alkaline RNA ladder (NaOH) obtained by partial hydrolysis in NaOH of the respective labeled RNAs and radioactive markers.

*Figure S3: AsflhD repression of the transcription of flhD is independent of the P_flhD_ promoter*

(A) Schematic representation of the template used for the *in vitro* transcription assay carrying the P*_tet-_flhD* promoter driving the expression of a 388 nts transcript (purple) and the P_AsflhD_ promoter driving the expression of a 260 nts transcript (orange). The DNA templates were constructed using the LM213 and LM9 oligonucleotides and carry chromosomal sequences starting at the *flhD* TSS with a 40 nts extension carrying the tetracycline promoter (78) so that the P_tet_ transcription starts at the position of the *flhD* TSS. (B) *In vitro* transcription assays were performed on templates carrying wt and P_AsflhD_^+^ and P_AsflhD_^-^ mutations and (C) on the wt template after addition of *in vitro* synthesized AsflhD or *flhD* transcripts to the reaction at the indicated concentrations (supplementary material and method). Relative intensity of the indicated bands (*flhD* in purple and AsflhD in orange) were quantified as described in the “quantification and statistical analysis” with a number of sample n=6 and n=3 (C). Values are means of 6 (B) or 3 (C) replicates and error bars are standard deviations. Statistical significance was determined by a heteroscedastic two-tailed t test (* for p-values ≤0.05, ** for p-values ≤0.01 and *** for p-values ≤0.001).

*Figure S4: AsflhD overexpression repress and delay the induction of the motility cascade*

MG1655-B (wt) and ML241 (P_AsflhD_^+^) were grown at 37°C until mid-log phase (A_600_ = 0.4) or until late-log phase (A_600_ = 1). Total RNA was analyzed by northern blotting. The membranes were probed successively for (A) *flgB* and M1 and for (B) *fliC* and M1.

## Bibliography

1. Keseler IM, Mackie A, Santos-Zavaleta A, Billington R, Bonavides-Martínez C, Caspi R, Fulcher C, Gama-Castro S, Kothari A, Krummenacker M, Latendresse M, Muñiz-Rascado L, Ong Q, Paley S, Peralta-Gil M, Subhraveti P, Velázquez-Ramírez DA, Weaver D, Collado-Vides J, Paulsen I, Karp PD. 2016. The EcoCyc database: reflecting new knowledge about *Escherichia coli* K-12. Nucleic Acids Res 45:543–550.

2. Theodorou MC, Theodorou EC, Kyriakidis DA. 2012. Involvement of AtoSC two-component system in *Escherichia coli* flagellar regulon. Amino Acids 43:833–844.

3. Lemke JJ, Durfee T, Gourse RL. 2009. DksA and ppGpp Directly Regulate Transcription of the *Escherichia coli* Flagellar Cascade. Mol Microbiol 74:1368–1379.

4. Mizushima T, Koyanagi R, Katayama T, Miki T, Sekimizu K. 1997. Decrease in expression of the master operon of flagellin synthesis in a dnaA46 mutant of *Escherichia coli*. Biol Pharm Bull 20:327–331.

5. De Lay N, Gottesman S. 2012. A complex network of small non-coding RNAs regulate motility in *Escherichia coli*. Mol Microbiol 86:524–538.

6. Yakhnin AV, Baker CS, Vakulskas CA, Yakhnin H, Berezin I, Romeo T, Babitzke P. 2013. CsrA activates *flhDC* expression by protecting *flhDC* mRNA from RNase E-mediated cleavage. Mol Microbiol 87:851–866.

7. Lejars M, Hajnsdorf E. 2020. The world of asRNAs in Gram-negative and Gram-positive bacteria. Biochim Biophys Acta Gen Reg Mech 1863.

8. Hör J, Matera G, Vogel J, Gottesman S, Storz G. 2020. Trans-Acting Small RNAs and Their Effects on Gene Expression in *Escherichia coli* and *Salmonella enterica*. EcoSal Plus doi:10.1128/ecosalplus.ESP-0030-2019.

9. Dornenburg JE, DeVita AM, Palumbo MJ, Wade JT. 2010. Widespread Antisense Transcription in *Escherichia coli*. mBio 1.

10. Wade JT, Grainger DC. 2014. Pervasive transcription: illuminating the dark matter of bacterial transcriptomes. Nat Rev Microbiol 12:647–653.

11. Lejars M, Kobayashi A, Hajnsdorf E. 2019. Physiological roles of antisense RNAs in prokaryotes. Biochimie 164:3–16.

12. Masachis S, Darfeuille F. 2018. Type I Toxin-Antitoxin Systems: Regulating Toxin Expression via Shine-Dalgarno Sequence Sequestration and Small RNA Binding. Microbiol Spectr 6.

13. Malmgren C, Wagner EGH, Ehresmann C, Ehresmann B, Romby P. 1997. Antisense RNA Control of Plasmid R1 Replication. The dominant product of the antisense RNA-mRNA binding is not a full RNA duplex. J Biol Chem 272:12508–12512.

14. Darfeuille F, Unoson C, Vogel J, Wagner EG. 2007. An antisense RNA inhibits translation by competing with standby ribosomes. Mol Cell 26:381–392.

15. André G, Even S, Putzer H, Burguière P, Croux C, Danchin A, Martin-Verstraete I, Soutourina O. 2008. S-box and T-box riboswitches and antisense RNA control a sulfur metabolic operon of *Clostridium acetobutylicum*. Nucleic Acids Res 36:5955–5969.

16. Giangrossi M, Prosseda G, Tran CN, Brandi A, Colonna B, Falconi M. 2010. A novel antisense RNA regulates at transcriptional level the virulence gene *icsA* of *Shigella flexneri*. Nucleic Acids Res 38:3362–3375.

17. Kolb FA, Malmgren C, Westhof E, Ehresmann C, Ehresmann B, Wagner EG, Romby P. 2000. An unusual structure formed by antisense-target RNA binding involves an extended kissing complex with a four-way junction and a side-by-side helical alignment. Rna 6:311–24.

18. Opdyke JA, Kang J-G, Storz G. 2004. GadY, a Small-RNA Regulator of Acid Response Genes in *Escherichia coli*. J Bacteriol 186:6698–6705.

19. Opdyke JA, Fozo EM, Hemm MR, Storz G. 2011. RNase III Participates in GadY-Dependent Cleavage of the *gadX*-*gadW* mRNA. J Mol Biol 406:29–43.

20. Peters JM, Mooney RA, Grass JA, Jessen ED, Tran F, Landick R. 2012. Rho and NusG suppress pervasive antisense transcription in *Escherichia coli*. Genes Dev 26:2621–2633.

21. Conway T, Creecy JP, Maddox SM, Grissom JE, Conkle TL, Shadid TM, Teramoto J, San Miguel P, Shimada T, Ishihama A, Mori H, Wanner BL. 2014. Unprecedented High-Resolution View of Bacterial Operon Architecture Revealed by RNA Sequencing. mBio 5.

22. Lybecker M, Zimmermann B, Bilusic I, Tukhtubaeva N, Schroeder R. 2014. The double-stranded transcriptome of *Escherichia coli*. Proc Natl Acad Sci U S A 111:3134–3139.

23. Thomason MK, Bischler T, Eisenbart SK, Forstner KU, Zhang A, Herbig A, Nieselt K, Sharma CM, Storz G. 2015. Global transcriptional start site mapping using differential RNA sequencing reveals novel antisense RNAs in *Escherichia coli*. J Bacteriol 197:18–28.

24. Huang L, Deighan P, Jin J, Li Y, Cheung H-C, Lee E, Mo SS, Hoover H, Abubucker S, Finkel N, McReynolds L, Hochschild A, Lieberman J. 2020. *Tombusvirus* p19 Captures RNase III-Cleaved Double-Stranded RNAs Formed by Overlapping Sense and Antisense Transcripts in *Escherichia coli*. mBio 11:e00485–20.

25. Maes A, Gracia C, Innocenti N, Zhang K, Aurell E, Hajnsdorf E. 2017. Landscape of RNA polyadenylation in *E. coli*. Nucleic Acids Res 45:2746–2756.

26. Stead MB, Marshburn S, Mohanty BK, Mitra J, Castillo LP, Ray D, van Bakel H, Hughes TR, Kushner SR. 2011. Analysis of *Escherichia coli* RNase E and RNase III activity *in vivo* using tiling microarrays. Nucleic Acids Res 39:3188–3203.

27. Hanamura A, Aiba H. 1991. Molecular mechanism of negative autoregulation of *Escherichia coli crp* gene. Nucleic Acids Res 19:4413–4419.

28. Okamoto K, Freundlich M. 1986. Mechanism for the autogenous control of the *crp* operon: transcriptional inhibition by a divergent RNA transcript. Proc Natl Acad Sci U S A 83:5000–5004.

29. Wurtzel ET, Chou MY, Inouye M. 1982. Osmoregulation of gene expression. I. DNA sequence of the ompR gene of the ompB operon of Escherichia coli and characterization of its gene product. J Biol Chem 257:13685–13691.

30. Kenney LJ, Anand GS. 2020. EnvZ/OmpR Two-Component Signaling: An Archetype System That Can Function Noncanonically. EcoSal Plus 9.

31. Pratt LA, Hsing W, Gibson KE, Silhavy TJ. 1996. From acids to osmZ: multiple factors influence synthesis of the OmpF and OmpC porins in *Escherichia coli*. Mol Microbiol 20:911–7.

32. Stincone A, Daudi N, Rahman AS, Antczak P, Henderson I, Cole J, Johnson MD, Lund P, Falciani F. 2011. A systems biology approach sheds new light on Escherichia coli acid resistance. Nucleic Acids Res 39:7512–28.

33. Zwir I, Shin D, Kato A, Nishino K, Latifi T, Solomon F, Hare JM, Huang H, Groisman EA. 2005. Dissecting the PhoP regulatory network of *Escherichia coli* and *Salmonella enterica*. Proceedings of the National Academy of Sciences of the United States of America 102:2862.

34. Groisman EA, Hollands K, Kriner MA, Lee EJ, Park SY, Pontes MH. 2013. Bacterial Mg2+ homeostasis, transport, and virulence. Annu Rev Genet 47:625–46.

35. Bertin P, Terao E, Lee EH, Lejeune P, Colson C, Danchin A, Collatz E. 1994. The H-NS protein is involved in the biogenesis of flagella in *Escherichia coli*. J Bacteriol 176:5537–5540.

36. Santos-Zavaleta A, Salgado H, Gama-Castro S, Sánchez-Pérez M, Gómez-Romero L, Ledezma-Tejeida D, García-Sotelo JS, Alquicira-Hernández K, Muñiz-Rascado LJ, Peña-Loredo P, Ishida-Gutiérrez C, Velázquez-Ramírez DA, Del Moral-Chávez V, Bonavides-Martínez C, Méndez-Cruz CF, Galagan J, Collado-Vides J. 2019. RegulonDB v 10.5: tackling challenges to unify classic and high throughput knowledge of gene regulation in *E. coli* K-12. Nucleic Acids Res 47:D212–d220.

37. Apirion D, Watson N. 1978. Ribonuclease III is involved in motility of *Escherichia coli*. J Bacteriol 133:1543–1545.

38. Mitchell JE, Zheng D, Busby SJW, Minchin SD. 2003. Identification and analysis of ‘extended –10’ promoters in *Escherichia coli*. Nucleic Acids Res 31:4689–4695.

39. Koo BM, Rhodius VA, Campbell EA, Gross CA. 2009. Dissection of recognition determinants of *Escherichia coli* sigma32 suggests a composite -10 region with an ‘extended -10’ motif and a core -10 element. Mol Microbiol 72:815–29.

40. Rhodius VA, Suh WC, Nonaka G, West J, Gross CA. 2005. Conserved and Variable Functions of the σE Stress Response in Related Genomes. PLOS Biol 4.

41. Thompson KM, Rhodius VA, Gottesman S. 2007. SigmaE regulates and is regulated by a small RNA in *Escherichia coli*. J Bacteriol 189:4243–4256.

42. Ades SE, Connolly LE, Alba BM, Gross CA. 1999. The Escherichia coli sigma(E)-dependent extracytoplasmic stress response is controlled by the regulated proteolysis of an anti-sigma factor. Genes Dev 13:2449–61.

43. Rouvière PE, De Las Peñas A, Mecsas J, Lu CZ, Rudd KE, Gross CA. 1995. *rpoE*, the gene encoding the second heat-shock sigma factor, sigma E, in *Escherichia coli*. EMBO J 14:1032–1042.

44. Gruber AR, Lorenz R, Bernhart SH, Neuböck R, Hofacker IL. 2008. The Vienna RNA websuite. Nucleic Acids Res 36:W70–4.

45. Soutourina O, Kolb A, Krin E, Laurent-Winter C, Rimsky S, Danchin A, Bertin P. 1999. Multiple control of flagellum biosynthesis in *Escherichia coli:* role of H-NS protein and the cyclic AMP-catabolite activator protein complex in transcription of the *flhDC* master operon. J Bacteriol 181:7500–7508.

46. Fitzgerald DM, Bonocora RP, Wade JT. 2014. Comprehensive mapping of the *Escherichia coli* flagellar regulatory network. PLoS Genet 10:e1004649.

47. Lehnen D, Blumer C, Polen T, Wackwitz B, Wendisch VF, Unden G. 2002. LrhA as a new transcriptional key regulator of flagella, motility and chemotaxis genes in *Escherichia coli*. Mol Microbiol 45:521–532.

48. Kim YJ, Im SY, Lee JO, Kim OB. 2016. Potential Swimming Motility Variation by AcrR in *Escherichia coli*. J Microbiol Biotechnol 26:1824–1828.

49. Kery MB, Feldman M, Livny J, Tjaden B. 2014. TargetRNA2: identifying targets of small regulatory RNAs in bacteria. Nucleic Acids Res 42:W124–W129.

50. Raghavan R, Sloan DB, Ochman H. 2012. Antisense Transcription Is Pervasive but Rarely Conserved in Enteric Bacteria. mBio 3.

51. Campos A, Matsumura P. 2001. Extensive alanine scanning reveals protein-protein and protein-DNA interaction surfaces in the global regulator FlhD from *Escherichia coli*. Mol Microbiol 39:581–594.

52. Albanna A, Sim M, Hoskisson PA, Gillespie C, Rao CV, Aldridge PD. 2018. Driving the expression of the *Salmonella enterica* sv Typhimurium flagellum using flhDC from *Escherichia coli* results in key regulatory and cellular differences. Sci Rep 8:16705.

53. Afonyushkin T, Večerek B, Moll I, Bläsi U, Kaberdin VR. 2005. Both RNase E and RNase III control the stability of *sodB* mRNA upon translational inhibition by the small regulatory RNA RyhB. Nucleic Acids Res 33:1678–1689.

54. Aiso T, Kamiya S, Yonezawa H, Gamou S. 2014. Overexpression of an antisense RNA, ArrS, increases the acid resistance of Escherichia coli. Microbiol 160:954–961.

55. Wei BL, Brun-Zinkernagel AM, Simecka JW, Pruss BM, Babitzke P, Romeo T. 2001. Positive regulation of motility and *flhDC* expression by the RNA-binding protein CsrA of *Escherichia coli*. Mol Microbiol 40:245–256.

56. Thomason MK, Fontaine F, De Lay N, Storz G. 2012. A small RNA that regulates motility and biofilm formation in response to changes in nutrient availability in *Escherichia coli*. Mol Microbiol 84:17–35.

57. Naville M, Gautheret D. 2009. Transcription attenuation in bacteria: theme and variations. Brief Funct Genomic Proteomic 8:482–92.

58. Sedlyarova N, Shamovsky I, Bharati BK, Epshtein V, Chen J, Gottesman S, Schroeder R, Nudler E. 2016. sRNA-Mediated Control of Transcription Termination in E. coli. Cell 167:111–121.e13.

59. Stork M, Di Lorenzo M, Welch TJ, Crosa JH. 2007. Transcription Termination within the Iron Transport-Biosynthesis Operon of *Vibrio anguillarum* Requires an Antisense RNA. J Bacteriol 189:3479–3488.

60. Brantl S, Birch-Hirschfeld E, Behnke D. 1993. RepR protein expression on plasmid pIP501 is controlled by an antisense RNA-mediated transcription attenuation mechanism. J Bacteriol 175:4052–4061.

61. Chatterjee A, Johnson CM, Shu C-C, Kaznessis YN, Ramkrishna D, Dunny GM, Hu W-S. 2011. Convergent transcription confers a bistable switch in *Enterococcus faecalis* conjugation. Proc Natl Acad Sci USA 108:9721–9726.

62. Sim M, Koirala S, Picton D, Strahl H, Hoskisson PA, Rao CV, Gillespie CS, Aldridge PD. 2017. Growth rate control of flagellar assembly in *Escherichia coli* strain RP437. Sci Rep 7.

63. Pruss BM, Matsumura P. 1997. Cell cycle regulation of flagellar genes. J Bacteriol 179:5602–5604.

64. Maeda K, Imae Y, Shioi JI, Oosawa F. 1976. Effect of temperature on motility and chemotaxis of *Escherichia coli*. J Bacteriol 127:1039–1046.

65. Rudenko I, Ni B, Glatter T, Sourjik V. 2019. Inefficient Secretion of Anti-sigma Factor FlgM Inhibits Bacterial Motility at High Temperature. iScience 16:145–154.

66. Lalaouna D, Massé E. 2015. Identification of sRNA interacting with a transcript of interest using MS2-affinity purification coupled with RNA sequencing (MAPS) technology. Genomics data 5:136–138.

67. Iosub IA, van Nues RW, McKellar SW, Nieken KJ, Marchioretto M, Sy B, Tree JJ, Viero G, Granneman S. 2020. Hfq CLASH uncovers sRNA-target interaction networks linked to nutrient availability adaptation. eLife 9:e54655.

68. Blattman SB, Jiang W, Oikonomou P, Tavazoie S. 2020. Prokaryotic single-cell RNA sequencing by *in situ* combinatorial indexing. Nat Microbiol doi:10.1038/s41564-020-0729-6.

69. Braun F, Hajnsdorf E, Regnier P. 1996. Polynucleotide phosphorylase is required for the rapid degradation of the RNase E-processed *rpsO* mRNA of *Escherichia coli* devoid of its 3’ hairpin. Mol Microbiol 19:997–1005.

70. Hajnsdorf E, Regnier P. 1999. *E. coli rpsO* mRNA decay: RNase E processing at the beginning of the coding sequence stimulates poly(A)-dependent degradation of the mRNA. J Mol Biol 286:1033–1043.

71. Hajnsdorf E, Carpousis AJ, Regnier P. 1994. Nucleolytic inactivation and degradation of the RNase III processed *pnp* message encoding polynucleotide phosphorylase of *Escherichia coli*. J Mol Biol 239:439–454.

72. Hajnsdorf E, Regnier P. 2000. Host factor Hfq of *Escherichia coli* stimulates elongation of poly(A) tails by poly(A) polymerase I. Proc Natl Acad Sci U S A 97:1501–1505.

73. Miller JH. 1972. Experiments in molecular genetics / Jeffrey H. Miller. Cold Spring Harbor Laboratory, New York.

74. Fontaine F, Gasiorowski E, Gracia C, Ballouche M, Caillet J, Marchais A, Hajnsdorf E. 2016. The small RNA SraG participates in PNPase homeostasis. RNA 22:1560–1573.

75. Folichon M, Arluison V, Pellegrini O, Huntzinger E, Regnier P, Hajnsdorf E. 2003. The poly(A) binding protein Hfq protects RNA from RNase E and exoribonucleolytic degradation. Nucleic Acids Res 31:7302–7310.

76. Maikova A, Peltier J, Boudry P, Hajnsdorf E, Kint N, Monot M, Poquet I, Martin-Verstraete I, Dupuy B, Soutourina O. 2018. Discovery of new type I toxin-antitoxin systems adjacent to CRISPR arrays in *Clostridium difficile*. Nucleic Acids Res 46:4733–4751.

77. Burgess RR. 2001. Sigma Factors, p 1831–1834. In Brenner S, Miller JH (ed), Encyclopedia of Genetics doi:https://doi.org/10.1006/rwgn.2001.1192. Academic Press, New York.

78. Coornaert A, Chiaruttini C, Springer M, Guillier M. 2013. Post-transcriptional control of the *Escherichia coli* PhoQ-PhoP two-component system by multiple sRNAs involves a novel pairing region of GcvB. PLoS Genet 9:e1003156.

